# The novel role of Kallistatin in linking metabolic syndromes and cognitive memory deterioration by inducing amyloid-β plaques accumulation and tau protein hyperphosphorylation

**DOI:** 10.1101/2024.05.20.594915

**Authors:** Weiwei Qi, Yanlan Long, Ziming Li, Zhen Zhao, Jinhui Shi, Wanting Xie, Laijian Wang, Yandan Tan, Ti Zhou, Minting Liang, Ping Jiang, Bin Jiang, Xia Yang, Guoquan Gao

## Abstract

Accumulation of amyloid β (Aβ) peptides and hyperphosphorylated tau proteins in the hippocampus triggers cognitive memory decline in Alzheimer’s disease (AD). The incidence and mortality of sporadic AD were tightly associated with diabetes and hyperlipidemia, while the exact linked molecular is uncertain. Here, we reported that serum Kallistatin concentrations were meaningfully higher in AD patients, with a higher concentration of fasting blood glucose and triglyceride. In addition, the constructed Kallistatin-transgenic (KAL-TG) mice defined its cognitive memory impairment phenotype and lower LTP in hippocampal CA1 neurons accompanied by increased Aβ deposition and tau phosphorylation. Mechanistically, Kallistatin could directly bind to the Notch1 receptor and thereby upregulate BACE1 expression by inhibiting PPARγ signaling, resulting in Aβ cleavage and production. Besides, Kallistatin could promote the phosphorylation of tau by activating GSK-3β. Fenofibrate, a hypolipidemic drug, could alleviate cognitive memory impairment by down-regulating Aβ and tau phosphorylation of KAL-TG mice. Collectively, our data clarified a novel mechanism for Aβ accumulation and tau protein hyperphosphorylation regulation by Kallistatin, which might play a crucial role in linking metabolic syndromes and cognitive memory deterioration, and suggested that fenofibrate might have the potential for treating metabolism-related AD.

**Highlights:**

- Kallistatin-transgenic (KAL-TG) mice defined its cognitive memory impairment phenotype accompanied by increased Aβ deposition and tau phosphorylation.
- Kallistatin could directly bind to the Notch1 receptor and thereby upregulate BACE1 expression by inhibiting PPARγ signaling.
- Fenofibrate could alleviate cognitive memory impairment and down-regulate the serum Kallistatin level.

## Introduction

Alzheimer’s disease (AD), the most prevalent irreversible neurodegenerative disorder associated with dementia in elderly individuals, is marked by a gradual decline in cognitive memory. Pathologically, AD is identified by the presence of extracellular amyloid-β (Aβ) plaques and intracellular neurofibrillary tangles (NFTs) ^1, 2, 3^. The Aβ cascade and tau protein hyperphosphorylation are the two primary hypotheses concerning AD. According to the Aβ cascade hypothesis, the excessive production of Aβ disrupts normal cellular functions, leading to synaptic dysfunction, neurodegeneration, tau hyperphosphorylation, and neuroinflammation, which ultimately result in memory impairment in individuals with AD and dementia ^4, 5^. Aβ peptides are derived from the sequential cleavage of amyloid precursor protein (APP) by β-secretase (β-site APP cleaving enzyme 1, BACE1) and γ-secretase, thus making this cleavage process significant in AD pathology ^6, 7^. BACE1 is considered a highly promising therapeutic target. Several potent BACE1 inhibitors have progressed to advanced stages in clinical trials, emphasizing the role of BACE1 in Aβ production ^8, 9, 10^. Tau, a microtubule-associated protein, naturally occurs in axons and regulates microtubule dynamics and axonal transport ^11^. In AD, tau undergoes a multistep transformation from a natively unfolded monomer to large aggregated forms, such as NFTs, another defining feature of AD ^12, 13^. Glycogen synthase kinase-3 (GSK3) is a key kinase involved in the initial steps of tau phosphorylation, with Wnt signaling being crucial in activating GSK-3β and GSK-3β-mediated tau phosphorylation ^14^. The physiological mechanisms underlying their interaction remain poorly understood.

There is a close relationship between metabolic disorders and cognitive impairment across the AD spectrum ^15, 16^. Nearly 95% of AD patients are categorized as sporadic patients, whose increasing incidence and mortality are strongly associated with type 2 diabetes mellitus (T2DM), obesity, and hyperlipidemia ^17, 18, 19^. About 37% of comorbidities between AD and diabetes have been reported in the Alzheimer’s Association Report ^20, 21^. As a result of the strong association and shared mechanism between AD and T2DM, AD has been termed “type 3 diabetes” by some researchers ^22, 23, 24, 25^. Several studies have demonstrated that diabetes confers a 1.6-fold increased risk of developing dementia ^26, 27^. Similarly, central obesity and high body mass index (BMI) during middle age are associated with an about 3.5-fold increased risk of dementia later in life ^28^. Therefore, controlling blood glucose and lipids is expected to be a strategy for preventing or moderating cognitive decline during aging. Nevertheless, the exact link and key associated regulators between metabolic abnormalities and AD are still unclear.

Kallistatin is a serine proteinase inhibitor previously identified as a tissue kallikrein-binding protein ^29^. It is produced predominantly in the liver and is widely expressed in body tissues, where it has antiangiogenic, antifibrotic, antioxidative stress, and antitumor effects ^30, 31^. Furthermore, Kallistatin was found to be increased in patients with obesity, prediabetes, and diabetes ^32, 33, 34^. The concentration of Kallistatin was positively correlated with the triglyceride glucose index ^35^, which was proven to be an independent risk factor for dementia ^36^. In addition, our previous study revealed that the concentration of serum Kallistatin in T2DM patients was significantly increased and further revealed that Kallistatin suppressed wound healing in T2DM patients by promoting local inflammation, which suggested that Kallistatin plays a critical role in the progression of T2DM ^37^. Furthermore, our recent research revealed that Kallistatin can cause memory and cognitive dysfunction by disrupting the glutamate-glutamine cycle^38^.

To explore the relationships among T2DM, AD and Kallistatin, we constructed Kallistatin transgenic (KAL-TG) mice to explore whether Kallistatin could cause cognitive impairment through the upregulation of Aβ production. Taken together, our results suggest that a novel regulatory mechanism of Aβ production and tau protein hyperphosphorylation by Kallistatin is involved in the progression of metabolic abnormality-related AD.

## Results

### Kallistatin increases in AD patients and AD model mice

To explore the relevance of AD in T2DM, a GAD disease enrichment analysis was initially conducted on differentially expressed genes in neurons of T2DM patients and normal controls, revealing a close relationship between AD and T2DM (GSE161355) (Fig. S1A). Additionally, PFAM analysis using the DAVID database identified enrichment of the Serpin family protein domain (Fig. S1B) (https://david.ncifcrf.gov/). Our previous studies revealed that Kallistatin (serpin family a member 4) was elevated in the serum of T2DM patients and was associated with an adverse prognosis of diabetes complications ^44^. We collected 11 serum samples from dementia patients at Sun Yat-sen Memorial Hospital and reported that the concentration of Kallistatin was greater than that in normal controls^38^. In this study, 56 AD patients and 61 healthy controls were enrolled from four hospitals in Guangdong Province to further investigate the potential relevance of Kallistatin and AD. The clinical and biochemical characteristics of the participants are provided in Tables S1 and S2. In addition, the serum Kallistatin (12.78 ±2.80 μg/mL) content in patients with AD was greater than that in normal controls (9.78 ±1.93 μg/mL) (Fig. 1A). Similarly, fasting blood glucose (FBG) and triglyceride (TG) levels were greater in AD patients than in healthy controls (Fig. 1B). We further grouped all the AD patients according to diabetes status and found that the Kallistatin (13.79 ±3.05 μg/mL) and TG contents were further elevated in AD patients with diabetes (Fig. 1C-D). Similarly, Kallistatin expression was increased in the hippocampus of the AD model mouse SAMP8 compared with that in the control mouse SAMR1 (Fig. S1C-D). Taken together, these results indicate that the Kallistatin concentration is increased in metabolic abnormality-related AD patients.

**Fig. 1.**
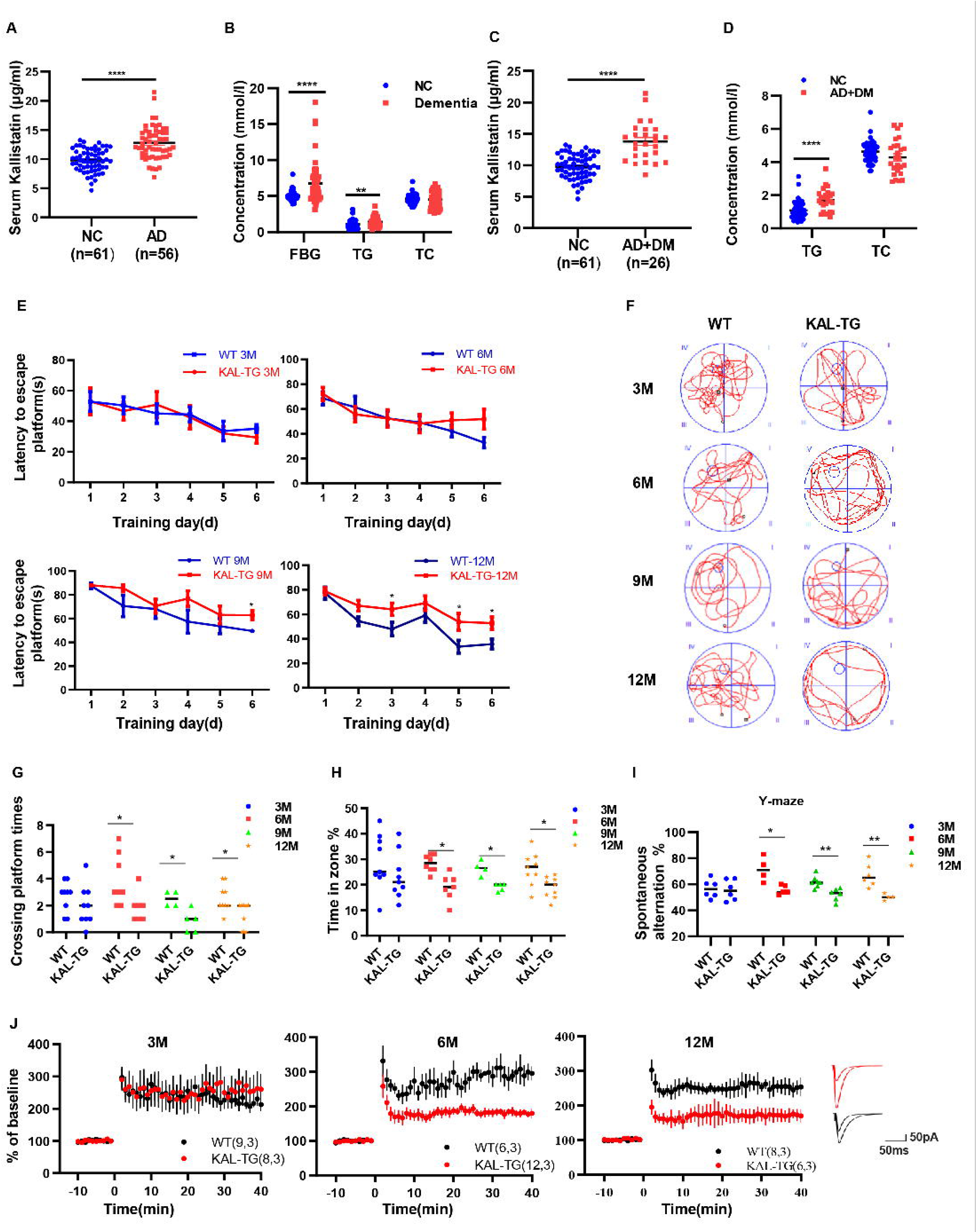
Increased Kallistatin was presented in AD patients and could impair cognitive memory in mice. (A-B) Serum Kallistatin(A), fasting blood glucose (FBG), triglyceride (TG), and total cholesterol (TC) (B) of AD patients and their corresponding normal control subjects. (C-D) Serum Kallistatin(C), TG, and TC(D) of AD patients with DM and their corresponding normal control subjects (Student’s t-test). (E-J) The behavioral performance of KAL-TG mice was assessed through the Morris water maze test, Y-maze test, and electrophysiology. (E)The escape latency time of different months of KAL-TG mice (3M, 6M, 9M, 12M) and corresponding WT mice were presented during 1-6 day (two-way ANOVA). (F-H) Cognitive functions were evaluated by spatial probe test at day 7 (Student’s *t*-test), the representative each group mice traces were shown (F), then analyzing each group mice crossing platform times (G) and time percent in the targeted area (H), n=4 to 9 per group. (I) Spontaneous alternation of Y-maze test. (J) LTP was measured by whole-cell voltage-clamp recordings of CA1 neurons in acute hippocampal slices of KAL-TG (3M, 6M, 12M) and WT mice (Student’s t-test, n=6-12 cells from 3 mice per group). Error bars represent the standard deviation (SD); one asterisk, *p* < 0.05, two asterisks, *p* < 0.01; four asterisks, *p* < 0.0001.

### Kallistatin impairs cognitive memory in mice

The above experiments demonstrated that Kallistatin was increased in AD patients and AD model mice. We subsequently generated KAL-TG mice and assessed their behavioral performance through the Morris water maze (MWM) and Y maze tests. Notably, the latency to escape the platform was longer, and the number of platform crossings, percentage of time spent, and spontaneous alternation were significantly lower in KAL-TG mice than in age-matched WT mice (Fig. 1E-I). Furthermore, long-term potentiation (LTP) was measured using whole-cell voltage-clamp recordings of CA1 neurons in acute hippocampal slices from KAL-TG and WT mice to assess changes in hippocampal synapses. The LTP in KAL-TG mice was significantly reduced compared to that in WT mice (Fig. 1J). These results showed that Kallistatin could impair cognitive memory in mice.

### Kallistatin promotes Aβ deposition and tau phosphorylation

We evaluated Aβ deposition and tau phosphorylation in these experimental mouse hippocampal tissues via immunohistochemistry (IHC) and western blotting. Predictably, the plaque density and tau phosphorylation in KAL-TG mice were much greater than those in age-matched WT mice (Fig. 2A-C, 3A-D). Consistent with these results, ELISA detection of the Aβ42 content in hippocampal tissue revealed that Aβ production was extraordinarily increased in KAL-TG mice compared with WT mice (Fig. 2D). These results suggested that Kallistatin promoted Aβ deposition and tau phosphorylation.

**Fig. 2.**
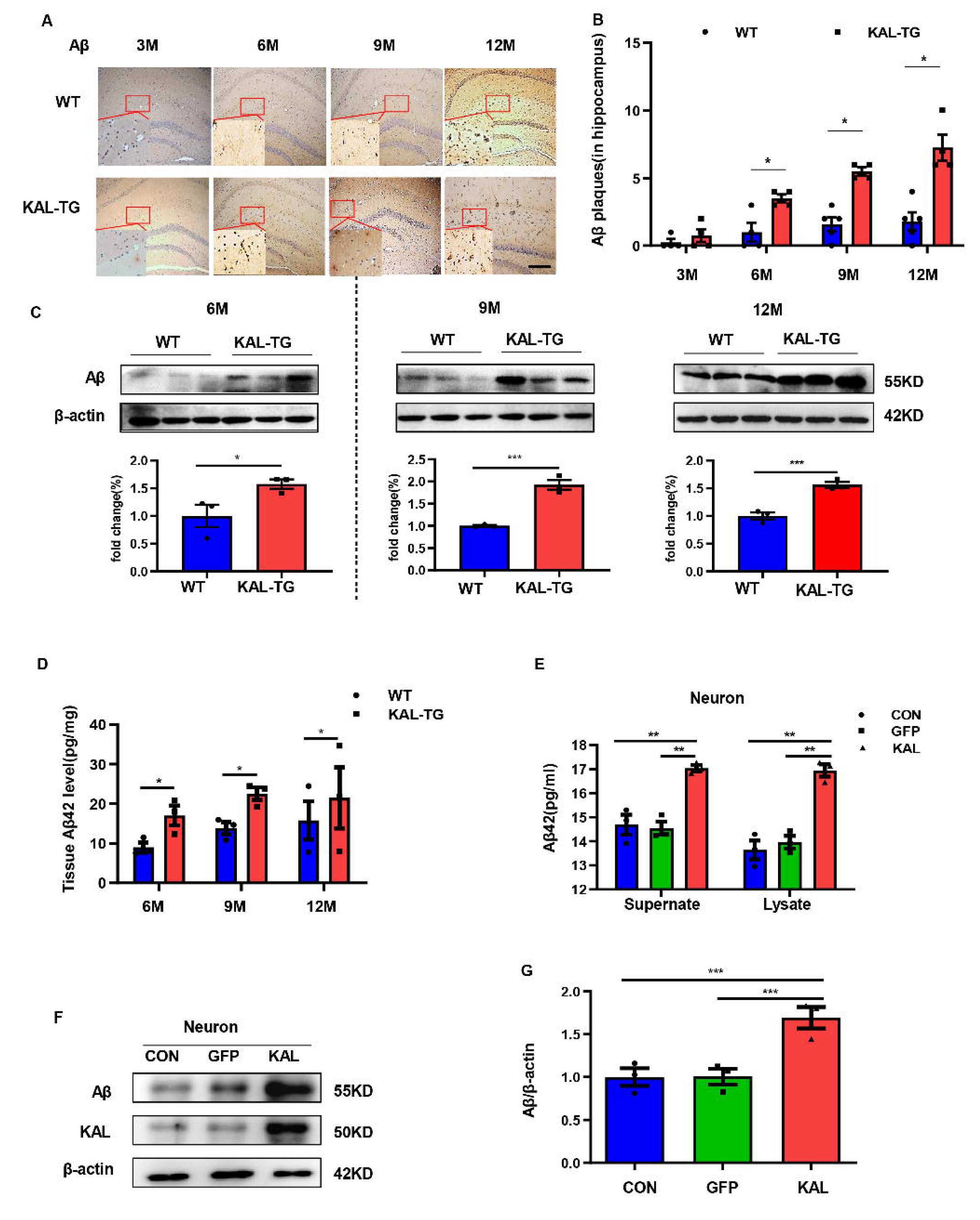
Kallistatin promoted Aβ generation. (A-B) Immunohistochemistry staining of Aβ(A) was carried out in KAL-TG and WT mice hippocampal tissue. *Scale bar*, 100μm. The statistical analysis of Aβ plaques (B) in hippocampal tissue of KAL-TG and WT mice, n=4-5 per group. (C) Protein levels of Aβ were tested by western blot analysis in hippocampal tissue, n=3 per group, then statistically analyzed the above results. (D) Hippocampal tissue Aβ42 contents were performed by ELISA in KAL-TG and WT groups, n=3 per group. (E) Primary mouse neurons were isolated, then infected with adenovirus to overexpress Kallistatin for 72h. Aβ42 concentration of primary hippocampal neurons supernate and cell lysate were quantified by ELISA, n=3 per group. (F-G) Western blot analysis of Aβ protein level in primary hippocampal neurons infected with overexpressing Kallistatin adenovirus and control groups, then statistical analysis of Aβ protein levels, n=3 per group. Error bars represent the standard deviation (SD); one asterisk, *p* < 0.05; two asterisks, *p* < 0.01; three asterisks, *p* < 0.001; Student’s *t*-test.

**Fig. 3.**
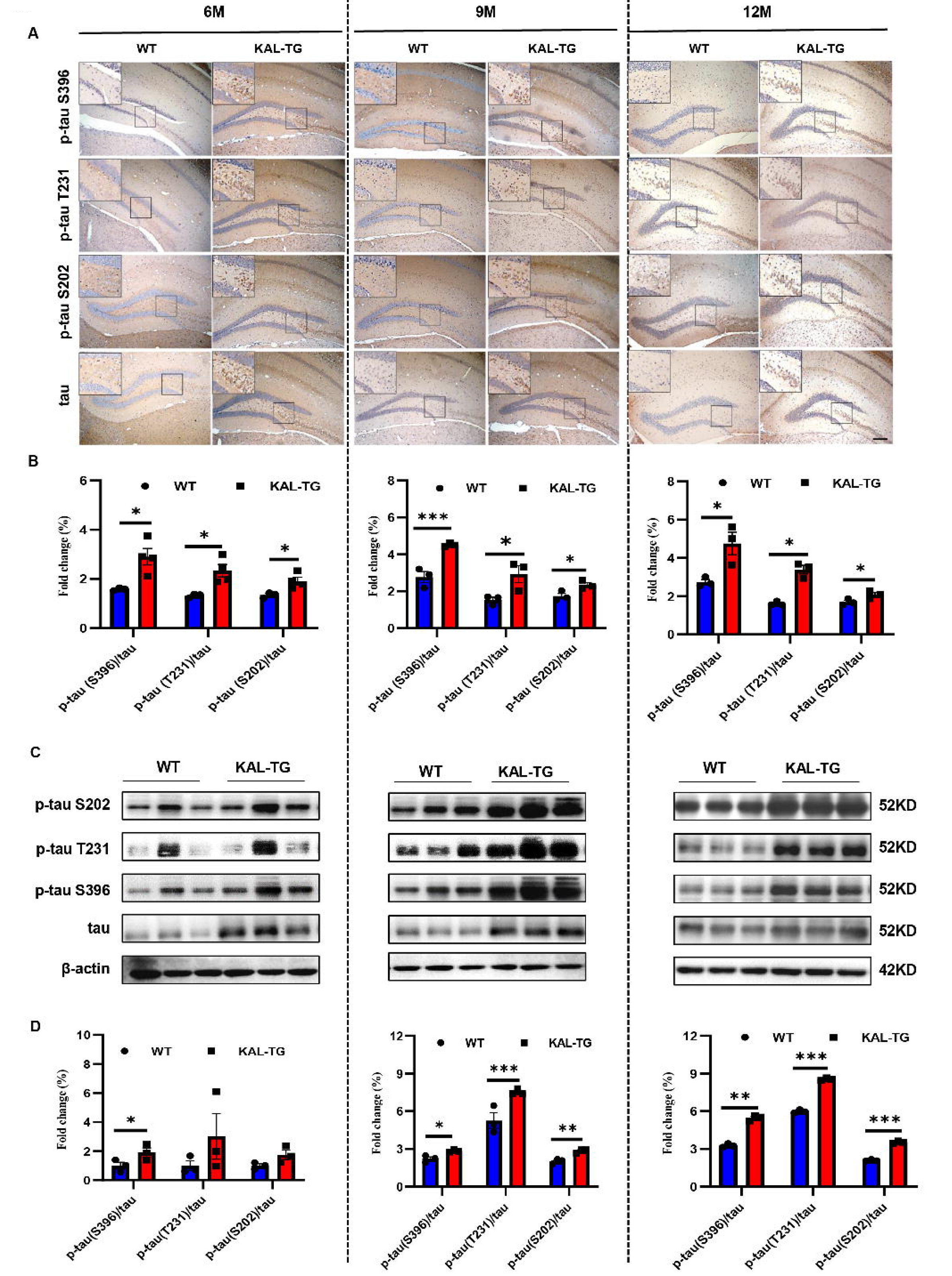
Kallistatin promoted tau phosphorylation. (A-B) Immunohistochemistry staining of phosphorylated tau (p-tau S396, p-tau T231, p-tau S202) and tau(A) was carried out in KAL-TG and WT mice hippocampal tissue. Scale bar, 100μm. The statistical analysis of phosphorylated tau(B) in hippocampal tissue of KAL-TG and WT mice, n=3 per group. (C-D) Protein levels of phosphorylated tau (p-tau S396, p-tau T231, p-tau S202) and tau were tested by western blot analysis in hippocampal tissue, then statistically analyzed the above results. Error bars represent the standard deviation (SD); one asterisk, p < 0.05; two asterisks, p < 0.01; three asterisks, p < 0.001; Student’s t-test.

### Kallistatin positively regulates Aβ generation by promoting β-secretase rather than γ-secretase

Western blot and ELISA analyses revealed that the Aβ levels in primary hippocampal neurons (immunofluorescence identified with the neural marker MAP2, Fig. S2G) infected with the Kallistatin adenovirus were greater than those in the control groups (Fig. 2E-G), as were those in the HT22 cells (Fig. S2A-C). Amyloid-beta precursor protein (APP) undergoes proteolytic processing to generate peptide fragments ^45^. β-Secretase (BACE1) and γ-secretase, which are composed of presenilin 1 (PS1), nicastrin, and Pen-2, are crucial enzymes for Aβ generation ^46, 47^. We determined the levels of APP, BACE1, and PS1 in hippocampal tissue. Compared with those in WT mice, BACE1 protein and mRNA levels were greater in KAL-TG mice (Fig. 4A-C, S2D), whereas no significant difference in APP, PS1, or α-secretase (ADAM9, ADAM10, or ADAM17) expression was detected (Fig. 4A, S2E). Consistent with the above results, the activity of BACE1 increased (Fig. 4D), whereas PS1 activity did not change (Fig. S2F). Similarly, the expression and activity of BACE1 were found to be increased in primary mouse neurons and HT22 cells transfected with Kallistatin adenovirus (Fig. 5A-C, S3A-C), while PS1 expression and activity remained unchanged (Fig. 5A, 5D, S3A). Additionally, the effect of Kallistatin was attenuated by the BACE1 inhibitor verubecestat or siBACE1 03, which was the most effective (Fig. 5E-F, S3D). These results indicate that Kallistatin can promote Aβ generation through the upregulation of BACE1 expression.

**Fig. 4.**
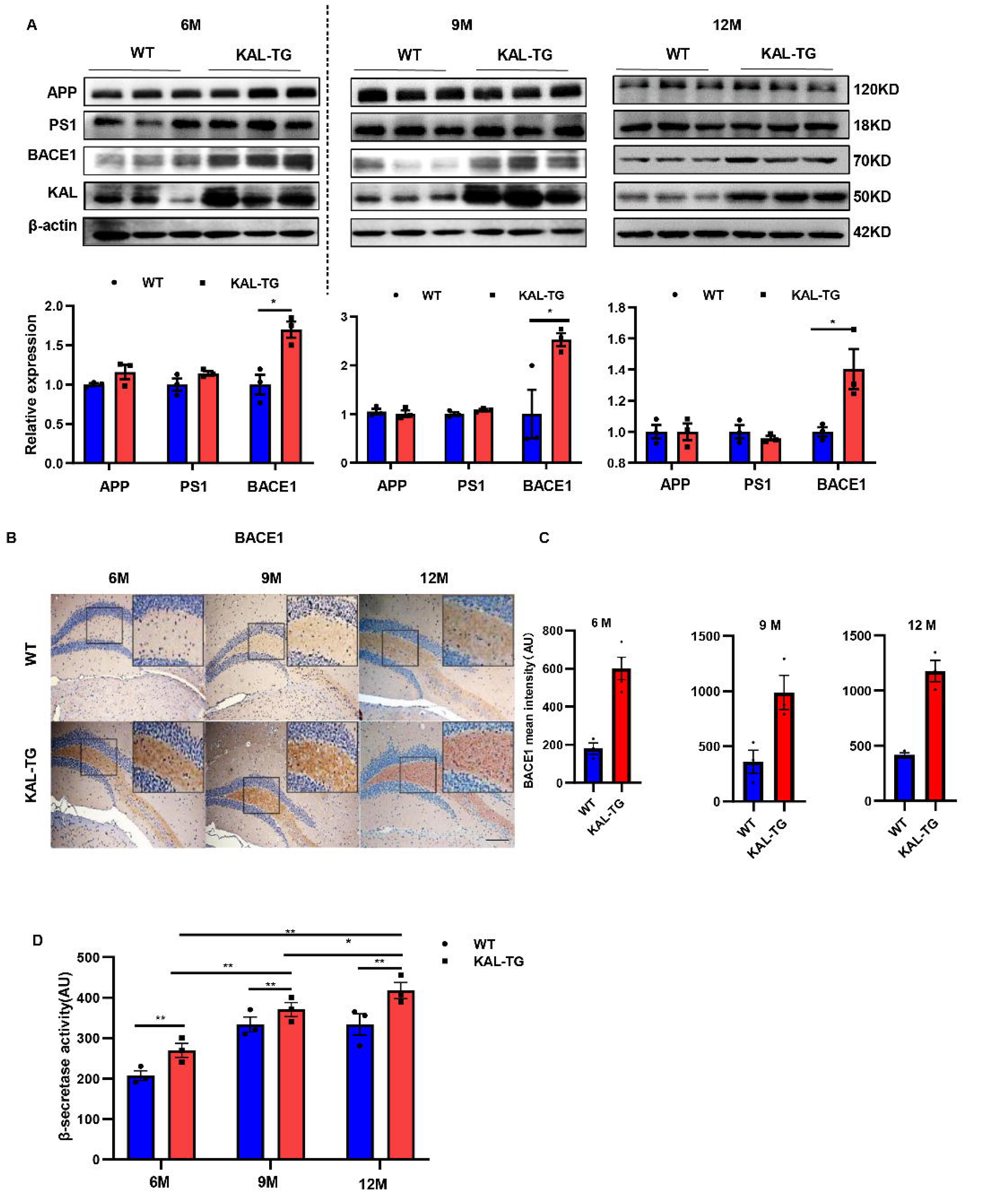
Kallistatin transgenic mice exhibited increased BACE1 expression and activity in the hippocampus. (A) Western blot analysis of relevant proteins, such as APP, PS1, and BACE1 during Aβ generation in hippocampal tissue of each time point (6M, 9M, 12M) KAL-TG mice and corresponding WT control groups, n=3 per group, then statistical analysis of APP, PS1 and BACE1 protein levels. (B) Immunohistochemistry staining of BACE1 was carried out in KAL-TG and WT mice hippocampal tissue at each time point (6M, 9M, 12M). n=3 to 5 per group. *Scale bar*, 100 μm. (C) Statistical analysis of BACE1 immunohistochemistry staining, n=3 to 4 per group. (D) ELISA measured the β-secretase activity of each group’s hippocampal tissue, n=3 per group. Error bars represent the standard deviation (SD); one asterisk, *p* < 0.05, two asterisks, *p* < 0.01; Student’s t-test.

**Fig. 5.**
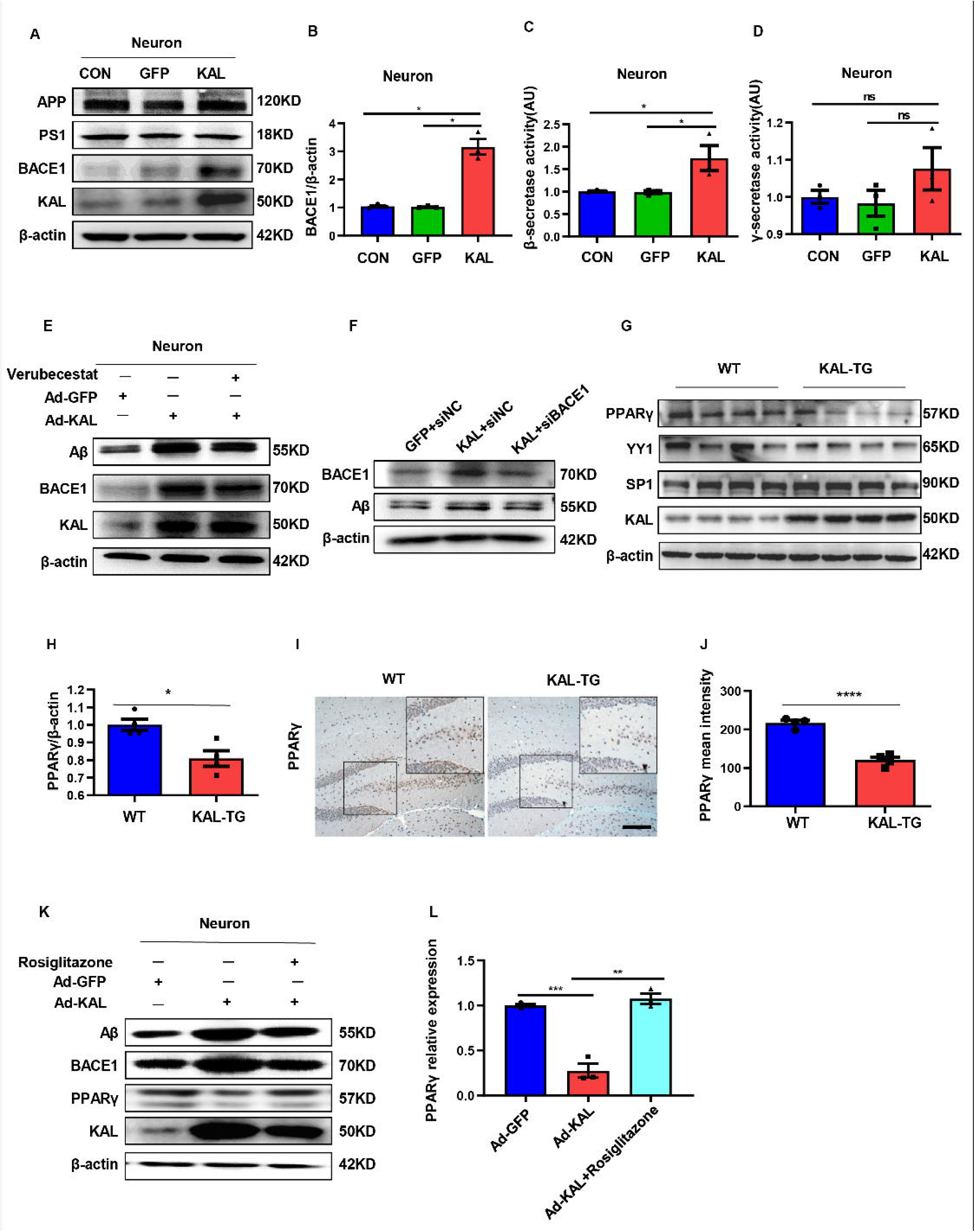
In *vitro*, Kallistatin promoted BACE1 expression to augment Aβ by suppressing PPARγ activation. (A) The relevant protein levels in primary mouse neurons infected with overexpressing Kallistatin adenovirus during Aβ generation were determined by western blot analysis. (B) Statistical analysis of BACE1 expression in primary neurons. (C-D) β-secretase(C) and γ-secretase(D) activity of primary hippocampal neurons infected with overexpressing Kallistatin adenovirus and control adenovirus was measured by ELISA. (E) Primary hippocampal neurons were treated with BACE1 inhibitor verubecestat (50nM), then infected with adenovirus to overexpress Kallistatin. Western blot analysis of Aβ, BACE1, and Kallistatin protein levels, β-actin served as a loading control. (F) HT22 cells were infected with BACE1 siRNA, then infected with adenovirus to overexpress Kallistatin. Western blot analysis of Aβ and BACE1 protein levels, β-actin served as a loading control. (G) The relevant proteins involved in BACE1 transcriptional expressions, such as PPARγ, YY1, and SP1 were measured by western blot analysis in hippocampal tissue. β-actin served as a loading control. (H) Statistical analysis of PPARγ in hippocampal tissue of each group. (I) The representative diagrams of PPARγ expression in hippocampal tissue were presented in the above graphs. *Scale bar*, 100μm. (J) Statistical analysis of PPARγ immunohistochemistry staining in hippocampal tissue of each group, n=3 to 4 per group. (K)Primary hippocampal neurons were treated with PPARγ agonist rosiglitazone (10nM) for 12h, then infected with adenovirus to overexpress Kallistatin for 72h. Western blot analysis of Aβ and BACE1 protein levels. β-actin served as a loading control. (L) Statistical analysis of PPARγ protein levels in each group. Error bars represent the standard deviation (SD), one asterisk, *p* < 0.05, two asterisks, *p* < 0.01; three asterisks, *p* < 0.001; ns means no significant difference; Student’s *t*-test.

### Kallistatin suppresses PPAR**γ** activation to promote BACE1 expression

The transcription factors SP1, YY1, and PPAR reportedly regulate BACE1 expression at the transcriptional level. Among them, PPARγ can downregulate BACE1 expression ^48, 49, 50^. PPARγ decreased in the hippocampal tissue of KAL-TG mice, as detected by western blot and immunohistochemical analysis (Fig. 5G-J). However, no significant differences in YY1 or SP1 expression were detected (Fig. 5G). In addition, Kallistatin downregulated the expression of PPARγ in primary hippocampal neurons and HT22 cells, thus increasing BACE1 and Aβ expression (Fig. 5K, S3E). Treatment with rosiglitazone, a PPARγ agonist, reversed the decrease in PPARγ caused by Kallistatin (Fig. 5K-L). Predictably, rosiglitazone inhibited the ability of Kallistatin to promote BACE1 and Aβ (Fig. 5K and Fig. S3E).

### Kallistatin promotes Aβ production *via* direct binding to the Notch1 receptor and activating the Notch1 pathway

Our results indicated that Notch1 was highly expressed in the hippocampal tissues of KAL-TG mice (Fig. 6A-D). Furthermore, Notch1 expression was upregulated in primary hippocampal neurons and HT22 cells infected with adenovirus expressing Kallistatin *in vitro* (Fig. S4A-B). Additionally, Kallistatin could directly bind to the Notch1 receptor and activate the Notch1 pathway (Fig. 6E-F and Fig. S4C-D). Treatment with siNotch1 03, which was the most effective (Fig. S4E), to knock down Notch1 inhibited the effect of Kallistatin on the activation of the Notch1 signaling pathway, leading to the downregulation of HES1, upregulation of PPARγ, and downregulation of BACE1 and Aβ (Fig. 6G). HES1, an essential downstream effector of the Notch1 signaling pathway, has been reported to suppress the expression of *PPARG* (the gene name of PPARγ) in neurons ^51, 52^. Similarly, HES1 shRNA 1, the most effective (Fig. S4F), upregulated PPARγ and decreased the production of BACE1 and Aβ when neurons were infected with adenovirus to overexpress Kallistatin (Fig. 6H). These results suggest that Kallistatin promotes Aβ production *via* direct binding to the Notch1 receptor and activating the Notch1 pathway.

**Fig. 6.**
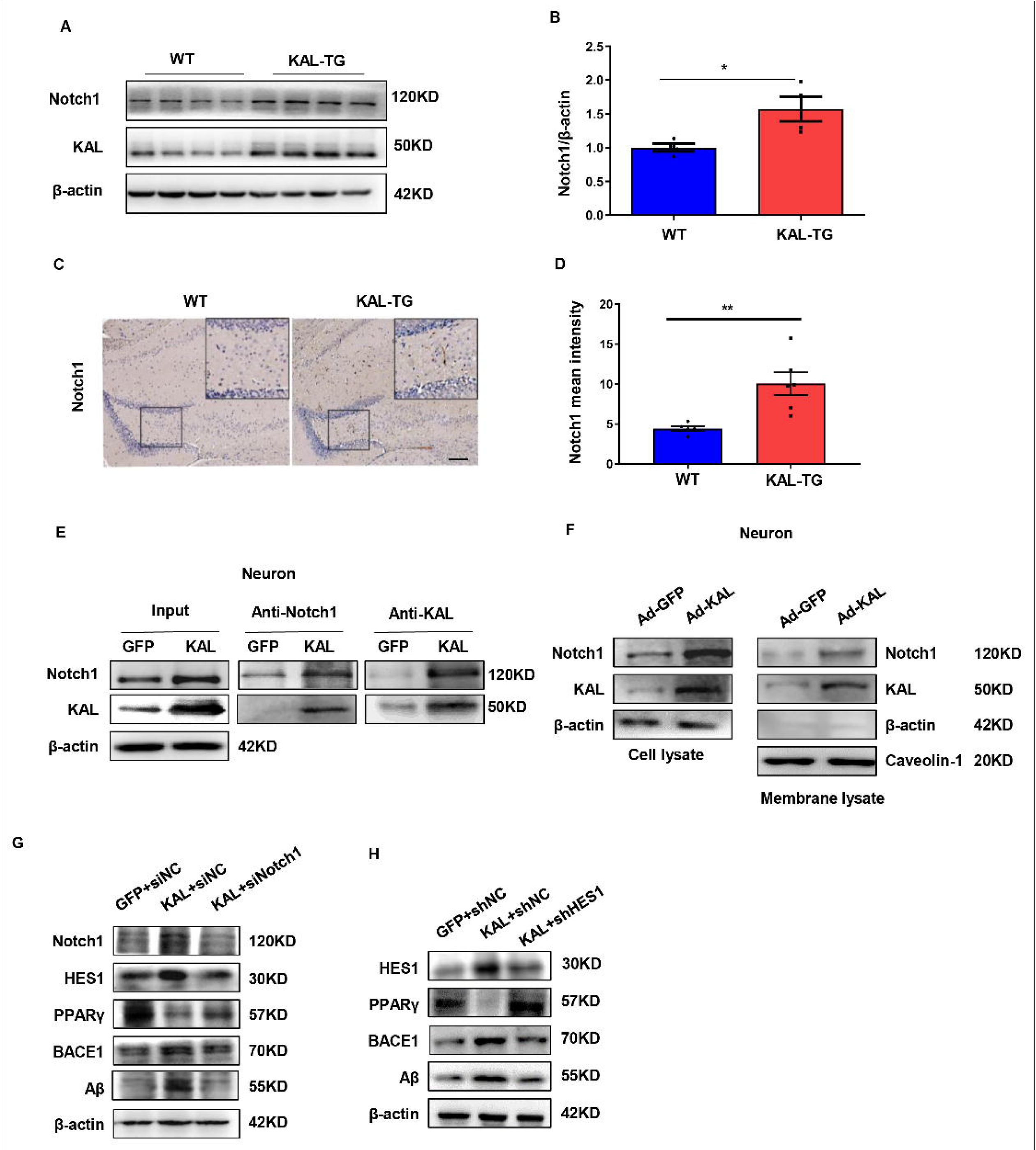
Kallistatin directly bonded to the Notch1 receptor, which activated the Notch1 pathway to promote Aβ production. (A) Notch1 expression was measured by western blot analysis in hippocampal tissue. β-actin served as a loading control. (B) Statistical analysis of Notch1 in hippocampal tissue of each group. (C) The representative diagrams of Nocth1 expression in hippocampal tissue were presented in the above graphs. Scale bar, 100μm. (D) Statistical analysis of Notch1 immunohistochemistry staining in hippocampal tissue of each group. (E-F) Primary hippocampal neurons were infected with overexpressing Kallistatin adenovirus for 72h, then Co-IP analysis (E) and membrane extraction experiment (F) was performed to verify whether Kallistatin can bind to the Notch1 receptor. β-actin served as a loading control. (G-H) HT22 cells were treated with siRNA (Notch1) and shRNA (HES1) to knock down Notch1 and HES1 for 12h, then infected with adenovirus to overexpress Kallistatin for 24h. Western blot analysis was used to detect the Notch1 signaling pathway. Error bars represent the standard deviation (SD), one asterisk, *p* < 0.05; Student’s t-test.

### Kallistatin promotes the phosphorylation of tau by activating the Wnt signaling pathway

Glycogen synthase kinase3-β (GSK-3β) is a crucial element in the phosphorylation of tau ^53^. When Wnt signaling is activated, the LRP6 PPPSPxS motif can directly interact with GSK-3β and phosphorylate it ^54^. Consequently, when Wnt signaling is inhibited, GSK-3β becomes activated and dephosphorylated, allowing nonphosphorylated GSK-3β to add phosphate groups to serine/threonine residues of tau ^14^. Kallistatin has already been reported as a competitive inhibitor of the canonical Wnt signaling pathway ^55^. Consistent with previous reports, our results demonstrated that GSK-3β was activated in the hippocampus of KAL-TG mice (Fig. 7A-B). Moreover, an increase in tau phosphorylation was observed with the activation of GSK-3β induced by Kallistatin overexpression (Fig. 7C-D, Fig. S5A), which was reversed by LiCl, an inhibitor of GSK-3β (Fig. 7E-F, Fig. S5B). These findings confirmed that Kallistatin promoted tau phosphorylation by activating the Wnt signaling pathway.

**Fig. 7.**
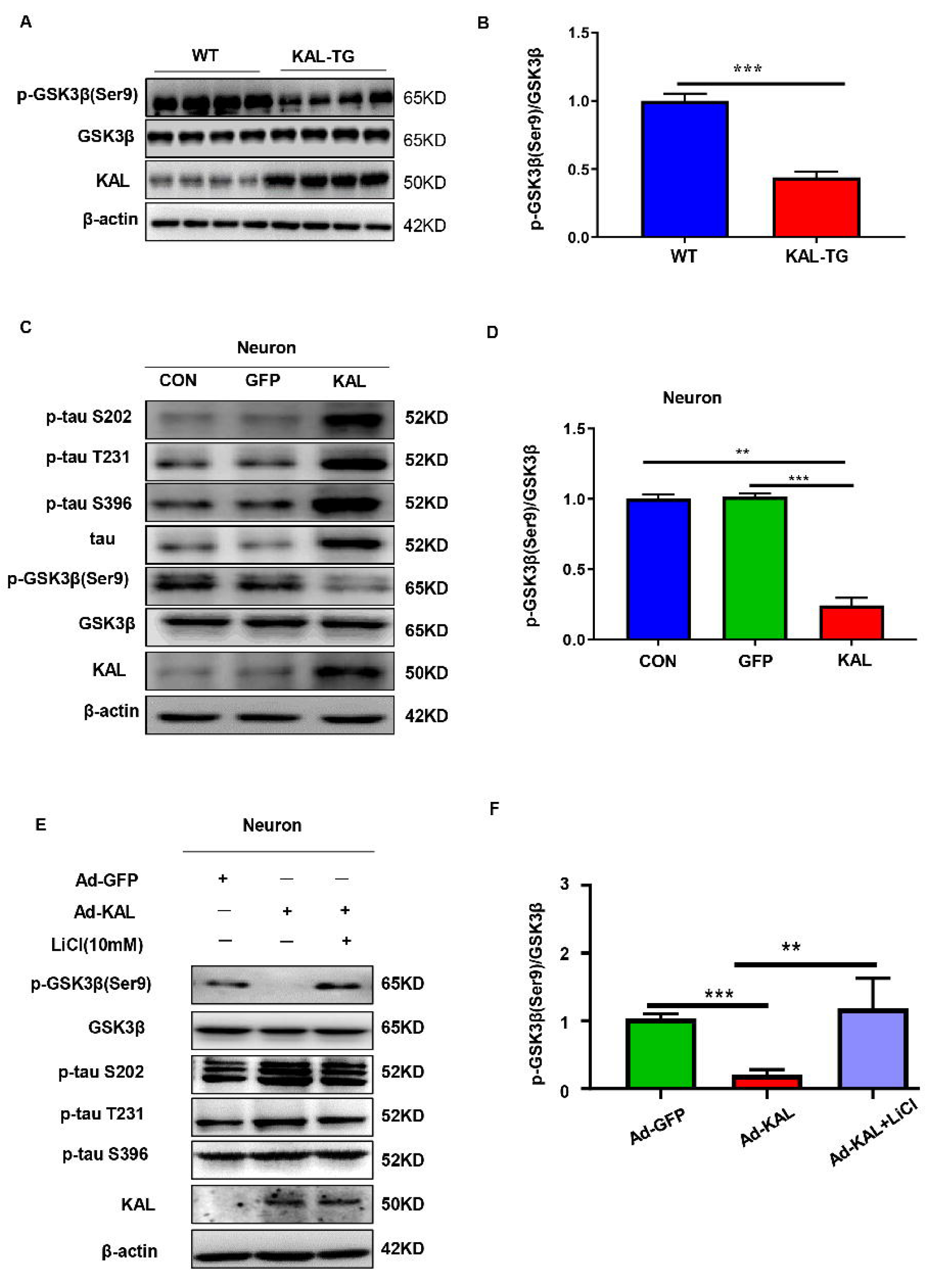
Kallistatin promoted phosphorylation of tau by suppressing Wnt signaling pathway. (A-B) GSK-3β and p-GSK-3β expression was measured by western blot analysis in hippocampal tissue, then statistically analyzed the above results. (C-D) Primary hippocampal neurons were treated with overexpressing Kallistatin adenovirus for 72h, then western blot analysis was used to detect the content of GSK-3β, p-GSK-3β, tau, p-tau (Ser9, T231, S396), and statistically analyzed the above results. (E-F) Primary hippocampal neurons were treated with overexpressing Kallistatin adenovirus for 48h, then treated with LiCl (10mM) for 24h, western blot analysis was used to detect the content of GSK-3β, p-GSK-3β, p-tau (Ser9, T231, S396), and statistical analysis of the above results. Error bars represent the standard deviation (SD), two asterisk, p < 0.01; three asterisk, p < 0.001; Student’s t-test.

### Fenofibrate alleviates memory and cognitive impairment in KAL-TG mice

Hyperlipidemia and hyperlipidemia account for the development of AD ^56, 57^. Here, a hypolipidemic drug (fenofibrate) and a hypoglycemic drug (rosiglitazone) were used to treat KAL-TG mice (Fig. 8A). Compared with that of the KAL-TG group, the behavioral performance of the treated group was improved, as measured by the MWM and Y-maze tests. The latency to reach the escape platform on the fifth training day was significantly decreased (Fig. 8B), and the number of platform crossings (Fig. 8C), percentage of time spent (Fig. 8D), and spontaneous alternation (Fig. 8F) were significantly greater in the fenofibrate-treated group than in the rosiglitazone-treated group. Similarly, the path trace heatmap indicated that the mice in the fenofibrate-treated group stayed in the target quadrant longer (Fig. 8E). In addition, decreased serum Kallistatin levels, Aβ and BACE1 levels, phosphorylation of tau, and activation of GSK3β were detected in the fenofibrate-treated KAL-TG group (Fig. 8G-K). However, there was no significant difference between the rosiglitazone-treated group and the KAL-TG group (Fig. 8G-K).

**Fig. 8.**
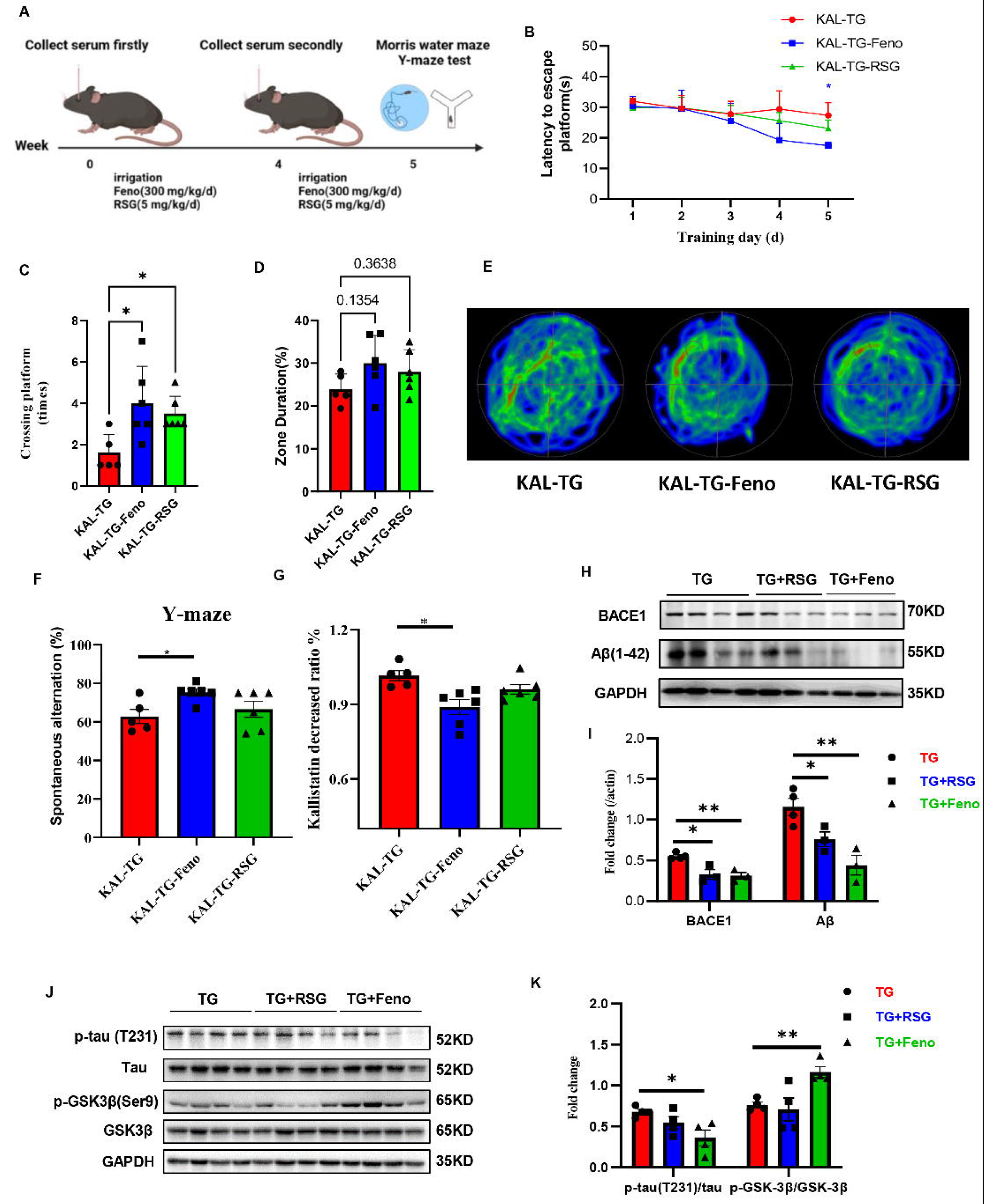
Fenofibrate could alleviate memory and cognitive impairment of KAL-TG mice. (A) Illustration of experimental protocols. Fenofibrate (0.3 g/kg/d × 5week, i.g.) or rosiglitazone (5mg/kg/d × 5week, i.g.) were given to KAL-TG mice. The serum for Kallistatin measuring was collected at week 0 and week 4. And Morris water maze and Y-maze test were performed at week 5. (B-E) Behavioral performance was assessed through the Morris water maze test and the Y-maze test. (B) The escape latency time was presented during 1-5 day. (C-E) Cognitive functions were evaluated by spatial probe test at day 6, then analyzing each group of mice crossing platform times(C), time percent in the targeted area (D), and the path traces heatmap (E), n=5 to 6 per group. (F) Spontaneous alternation of Y-maze test. (G) Kallistatin decreased ratio was calculated by dividing the serum Kallistatin concentration of KAL-TG mice before Fenofibrate/ rosiglitazone treatment by the serum Kallistatin concentration of KAL-TG mice after a-month treatment, and serum Kallistatin concentration was measured by ELISA. (H-I) Protein levels of Aβ and BACE1 were tested by western blot analysis in hippocampal tissue, then statistically analyzing the above results. (J-K) Protein levels of p-tau(231), tau, p-GSK-3β(Ser9) and GSK-3β were tested by western blot analysis in hippocampal tissue, then statistically analyzing the above results. Error bars represent the standard deviation (SD); one asterisk, *p* < 0.05; Student’s t-test.

### Mechanism summary

In patients with metabolic abnormality-related AD, the concentration of Kallistatin is elevated, which could increase Aβ deposition through the Notch1/HES1/PPARγ/BACE1 pathway and induce tau hyperphosphorylation by activating GSK-3β. Consequently, elevated Kallistatin impaired cognitive memory by inducing Aβ deposition and tau hyperphosphorylation (Fig. S5C).

## Discussion

This study demonstrated that Kallistatin is a novel regulator of amyloid-β plaque accumulation, tau protein hyperphosphorylation, and metabolic abnormality-related cognitive memory impairment. It was shown that Kallistatin levels were increased in the serum of patients with AD and diabetes-related AD, as well as in the hippocampus of AD model mice. Additionally, the KAL-TG mice exhibited cognitive memory impairment and lower LTP in hippocampal CA1 neurons, along with increased Aβ deposition and tau phosphorylation. Mechanistically, Kallistatin transcriptionally upregulates BACE1 expression by suppressing the transcriptional repressor PPARγ, leading to Aβ cleavage and production. Most importantly, our studies revealed that Kallistatin could bind directly to the Notch1 receptor and activate the Notch1/HES1 pathway, causing a decrease in PPARγ, overproduction of BACE1, and increased Aβ42 generation. Moreover, Kallistatin can induce tau phosphorylation by activating GSK-3β, which results from the inhibition of LRP6. Finally, prolonged stimulation with high concentrations of Kallistatin impaired cognitive memory in mice. Finally, the hypolipidemic drug fenofibrate decreased Aβ expression, tau phosphorylation, and the serum Kallistatin level in KAL-TG mice, alleviating memory and cognitive impairment. For the first time, these observations establish an association between high Kallistatin levels and metabolic abnormality-related AD and provide a new drug candidate (fenofibrate) for AD patients with metabolic syndromes.

Increasing evidence suggests that diabetes mellitus and AD are closely linked. About 80% of AD patients are insulin resistant or have T2DM ^58^; additionally, T2DM patients have a higher risk of up to 73% dementia than healthy controls do ^26^. In line with these observations, the process of cognitive decline in T2DM patients appears to begin in the prediabetic phase of insulin resistance ^59, 60^. A GAD disease enrichment analysis of differentially expressed genes in the neurons of T2DM patients and normal controls revealed a close relationship between AD and T2DM. Additionally, PFAM analysis identified an enrichment of the Serpin family protein domain (Fig. S1A, B). We and other researchers reported that the level of Kallistatin (which belongs to the serpin family) was increased in T2DM patients ^32, 37^. Although the relationship between Kallistatin and AD has not been reported to date, we speculate that Kallistatin might be a critical molecule that could establish a relationship between Alzheimer’s disease and diabetes. Our data indicated that AD patients exhibit metabolic disorders and elevated Kallistatin levels (Fig. 1A-D, S1C-D). Additionally, KAL-TG mice showed impaired memory, cognitive function, and synaptic plasticity (Fig. 1E-J). These results suggest that Kallistatin represents a novel connection between AD and diabetes.

A hallmark of AD is the aggregation of Aβ into amyloid plaques and tau phosphorylation in patients’ brains. Aβ, a small peptide with a high propensity to form aggregates, is widely believed to be central and initial to the pathogenesis of this disease ^61^. Correspondingly, it was discovered that Kallistatin could lead to Aβ overproduction through the Notch1/HES1/PPARγ/BACE1 signaling pathway. GSK-3β, a vital kinase that regulates the process of tau phosphorylation ^53^, can be activated by inhibiting the Wnt signaling pathway. Kallistatin has been identified as a competitive inhibitor of LRP6, the Wnt receptor ^55^. Consistent with previous reports, Kallistatin increased tau phosphorylation by activating GSK-3β. It was demonstrated for the first time that Kallistatin promoted AD by increasing Aβ production and tau phosphorylation in the central nervous system.

Previous studies have shown that Notch signaling is closely related to AD. For example, a NOTCH mutation was reported to cause AD-like pathology ^62^. In addition, the level of Notch1 was found to be increased in AD patients ^63^. In addition, the Notch1/HES1 signaling pathway has been reported to suppress the expression of PPARγ ^64^. Our previous studies demonstrated that Kallistatin can activate Notch1 signaling ^37^. Consequently, we detected Notch1 in our animal and cell models. Indeed, Notch1 was upregulated by Kallistatin (Fig. 6, S4), as was Aβ deposition. Notch signaling is initiated by receptor□ligand interactions at the cell surface. In mammals, there are five ligands encoded by JAG1, JAG2, DLL1, DLL3, and DLL4 ^65^. Here, our results revealed that Kallistatin could activate Notch1 by binding directly to it, indicating that Kallistatin is a new ligand of the Notch1 receptor.

The treatment of AD has always been a prominent and challenging issue in neurology. Multiple strategies have been proposed to reduce the pathogenicity of Aβ and tau. Unfortunately, several Aβ-targeted therapies tested in phase III clinical trials have failed to slow cognitive decline, although they can effectively reduce the Aβ load ^66, 67, 68, 69, 70, 71^. BACE1 inhibitors have not only failed to improve the cognitive function of AD patients but have also resulted in clinical deterioration and liver function impairment ^72, 73, 74^. Two possible reasons for the failure of clinical trials with BACE1 inhibitors are: first, the reduction in BACE1 activity could lead to the accumulation of full-length APP ^75^; second, the size of the BACE1 active site is relatively large, and the use of a small molecule may not be sufficient to occupy the active site ^76^. Consequently, synaptic damage caused by BACE1 inhibitors or their insufficient effect may lead to the failure of clinical trials. Therapeutic strategies targeting tau include tau aggregation blockers (TRx0014, TRx0237), antibody vaccine therapy (e.g., RO7105705, BIIB092), the inhibition of tau phosphorylation (Anavex2-73), and the use of microtubule stabilizers (Anavex2-73) ^77^. Some of these drugs have been partially discontinued, while others are still undergoing clinical testing and have shown protective benefits. Nonetheless, several obstacles remain to the commercialization of tau treatments when they reach maturity.

Because of the failure of clinical trials, some researchers have proposed alternative options for AD therapeutics to address modifiable risk factors for the development of AD, such as type 2 diabetes ^78, 79, 80^. Previous studies revealed that Kallistatin is a multifunctional protein strongly associated with diabetes and that a Kallistatin neutralizing antibody improves diabetic wound healing ^44^. This study demonstrated that Kallistatin induced memory-related cognitive dysfunction by promoting Aβ deposition and tau phosphorylation. Thus, it is speculated that increased Kallistatin could be a promising candidate for T2DM-related AD therapy. PPARγ is a ligand-activated transcription factor and a master modulator of glucose and lipid metabolism, organelle differentiation, and inflammation ^81, 82^. Growing evidence revealed that PPARγ agonists (rosiglitazone) could rescue memory impairment of AD model mice ^83, 84, 85^. In clinical trials, it is controversial whether rosiglitazone has a protective effect on memory cognitive function ^86, 87, 88^. In this study, although Aβ expression had a downward trend, the memory and cognition of KAL-TG mice were unchanged after treatment with rosiglitazone for a month (Fig.8). This might be caused by insufficient treatment time and the unchanged Kallistatin level.

Fenofibrate is a fibric acid derivative for clinically lowering blood lipids, mainly triglycerides ^89^. Studies showed that fenofibrate could prevent memory disturbances, maintain hippocampal neurogenesis, and protect against Parkinson’s disease (PD) ^90, 91^. Specifically, fenofibrate has a neuroprotective effect on memory impairment induced by Aβ through targeting α- and β-secretase ^92^. Recently, our study proved that the fenofibrate could repair the disrupted glutamine-glutamate cycle by upregulating glutamine synthetase, while there is currently no fenofibrate treatment of AD in clinical trials^38^. In this study, we proved fenofibrate was beneficial for memory and cognitive impairment of KAL-TG mice. In addition, Aβ, BACE1, phosphorylated tau, and serum Kallistatin level of KAL-TG mice could be downregulated after the treatment of fenofibrate. All of these suggested that fenofibrate might be helpful for metabolic abnormalities-related AD patients. Therefore, fenofibrate administration in patients with metabolic syndrome play an early role in preventing and treating AD.

In summary, we affirmed that Kallistatin concentrations were increased in diabetes-related AD patients. In addition, our study demonstrated for the first time that Kallistatin positively regulated Aβ42 through Notch1/HES1/PPARγ/BACE1, and increased phosphorylated tau through inhibition of the Wnt signaling pathway. Kallistatin might play a crucial role in linking diabetes and cognitive memory deterioration. Moreover, fenofibrate could decrease the serum Kallistatin level, BACE1, Aβ, and phosphorylated tau of KAL-TG mice, leading to alleviating memory and cognitive impairment. These findings might provide new insight into AD and possibly other neurodegenerative disorders.

## Materials and methods

### Ethics approval and consent to participate

All patients involved in this study gave their informed consent. This study obtained the institutional review board approval of Medical Ethics of Zhongshan Medical College No. 072 in 2021 and Animal Experiment Ethics of Zhongshan Medical College, Approval No.: SYSU-IACUC-2019-B051.

### Human Subjects

The study was approved by the experimental ethics committee of Guangdong Academy of Medical Sciences and Sun Yat-sen University and carried out in strict accordance with the ethical principles, and each participant was provided written informed consent before collecting samples. We certify that the study was performed in accordance with the 1964 declaration of HELSINKI and later amendments. We collected 61 normal human samples, 56 AD patient samples, of whom 36 normal human samples were from the Zhongshan City People’s Hospital; 14 normal human samples and 22 AD patient samples were from Zhongshan Third People’s Hospital; 11 normal human samples and 14 AD patient samples were from Sun Yat-sen Memorial Hospital. AD patient was clinically diagnosed according to ICD-10 (International Classification of Diseases) and NINCDS-ADRDA (the National Institute of Neurological and Communicative Disorders and Stroke and the Alzheimer’s Disease and Related Disorders Association) criteria, and 20 AD patient samples were clinically diagnosed according to MMSE (Mini-Mental State Examination), were collected from Guangdong Provincial People’s Hospital. All subjects’ Clinical characteristics were presented in Supplementary Tables (Table S1-2).

### Experimental Animals and Protocols

All animal experiment procedures were carried out in an environment without specific pathogens (Specific pathogen free, SPF) with the approval of the Animal Care and Use Committee of Sun Yat-sen University (approval ID: SYXK 2015-0107). The wild type mice (WT, C57BL/6) were purchased from the Animal Center of Guangdong Province (Production license No.: SCXK 2013-0002, Guangzhou, China). The SAMR1 and SAMP8 mice (7 months old) were purchased from Tianjin University of Traditional Chinese Medicine (Tianjin, China). Kallistatin transgenic mice (KAL-TG) were C57BL/6 strain provided by Dr. Jianxing Ma (University of Oklahoma Health Sciences Center) ^39^. The KAL-TG mice genotype was identified by PCR technology (forward primer: 5’-AGGGAAG-ATTGTGGATTTGG-3’, reverse primer: 5’-ATGAAGATACCAGTGATGCTC-3’). KAL-TG mice aged 6 months were randomly divided into three groups: control group (KAL-TG), fenofibrate-treated group (KAL-TG-Feno, 0.3□g/kg/d), and rosiglitazone-treated group (KAL-TG-RSG, 0.005□ g/kg/d). Fenofibrate (Sigma-Aldrich, cat. no. F6020) and rosiglitazone (Selleck, cat. no. S2556) were administered to mice by oral gavage. In three groups, the serum Kallistatin were examined in the 0 week and 4 week after drug treatment from the blood taken from mouse orbit. In addition, the Morris water maze and Y-maze test were performed one week after the second blood collection.

### Morris water maze (MWM)

The KAL-TG and WT mice were employed for the Morris water maze test including the behavioral test, latency experiment (for 6 days), and the probe test (the 7th day). In addition, the MWM was performed as described previously ^40^. Mice were brought into the testing room and handled for 1 day before the training experiment. In the 6-day training experiment, each mouse was trained with four daily trials. The mice facing the wall were placed into the maze, exploring the maze from different directions (east, south, west, and north). This trial was completed as soon as the mouse found the platform, or 90 s elapsed. If the mice could discover and climb the submerged platform within 90 s, the system would automatically record the latency time and path immediately, and then the mouse was guided to and placed on the submerged platform for extra 20 s. On day 7, the platform was removed, and a probe test was performed to examine the strength and integrity of the animal spatial memory 24 h after the last testing trial. During the probe test, the mice were gently brought into the water from the fixed monitoring point, and the mice were allowed to swim for 90 s without the platform. Finally, all of the measured behavioral parameters were analyzed using SMART software.

### Y-maze test

A Y maze test was performed to assess the mice’s spatial memory. The Y maze was separated by 120°, consisting of three identical arms (30 cm long, 7 cm wide, and 15 cm high) made of blue PVC. The mice were placed first in one of the arms, and over the next 10 minutes, the sequence and number of their entry into the three arms were monitored. An alternation is defined when a mouse visits three straight arms (namely, ABC, BCA, or CAB, but not ABA, BAB, or CAC). Spontaneous alternation (%)□=□[(number of alternations)/(total number of arms−2)]×□100.

### Electrophysiology

Hippocampal slices (300-400 μm) from KAL-TG and WT mice were cut as described ^41^. Coronal slices from hippocampus (400 μm thick) were prepared from different age groups KAL-TG mice and their WT littermates using a tissue slicer (Vibratome 3000; Vibratome) in ice-cold dissection buffer containing the following (in mM): 212.7 sucrose, 3 KCl, 1.25 NaH_2_PO_4_, 3 MgCl_2_, 1 CaCl_2_, 26 NaHCO_3_, and 10 dextrose, bubbled with 95% O_2_/5% CO_2_. The slices were immediately transferred to ACSF at 35 °C for 30 min before recordings. The recipe of ACSF was similar to the dissection buffer, except that sucrose was replaced with 124 mM NaCl, and the concentrations of MgCl_2_ and CaCl_2_ were changed to 1 mM and 2 mM, respectively. All recordings were performed at 28-30 °C. Pyramidal cells in CA1 areas were identified visually under infrared differential interference contrast optics based on their pyramidal somata and prominent apical dendrites. Whole-cell was performed using an Integrated Patch-Clamp Amplifier (Sutter Instrument, Novato, CA, USA) controlled by Igor 7 software (WaveMetrics, Portland, OR, USA) filtered at 5□kHz and sampled at 20□kHz. Igor 7 software was also used for acquisition and analysis. Only cells with series resistance <20 MΩ and input resistance >100 MΩ were studied. Cells were excluded if input resistance changed >15% or series resistance changed >10% over the experiment. A concentric bipolar stimulating electrode with a tip diameter of 125 μm (FHC) was placed in the stratum radiatum. The recording and stimulating electrode distances were kept at 50-100 μm. Patch pipettes (2-4MΩ) were filled with the internal solution consisting of the following (in mM): 120 Cs-methylsulfonate, 10 Na-phosphocreatine, 10 HEPES, 4 ATP, 5 lidocaine N-ethyl bromide (QX-314), 0.5 GTP; the pH of the solution was 7.2–7.3, and the osmolarity was 270-285 mOsm.

To induce LTP, a pairing protocol was applied. In brief, conditioning stimulation consisted of 360 pulses at 2 Hz paired with continuous postsynaptic depolarization (180 s) to 0 mV. 50 μM picrotoxin was added to the recording bath to suppress excessive polysynaptic activity, and the concentration of Ca^2+^ and Mg^2+^ was elevated to 4 mM to reduce the recruitment of polysynaptic responses. A test pulse was delivered at 0.067 Hz to monitor baseline amplitude for 10 min before and 30 min following paired stimulation. To calculate LTP, the EPSC amplitude was normalized to the mean baseline amplitude during 10 min baseline. Potentiation was defined as the mean normalized EPSC amplitude 25–40 min after paired stimulation.

### ELISA

To quantify serum Kallistatin, the collected samples were centrifuged at 4 □ for 10min at 5000 rpm. It was detected using the KBP ELISA kit (#DY1669, R&D systems, MN, USA) as per the instructions of the manufacturer. The levels of Aβ42 in brain tissue produced from mouse primary neuron cells and HT22 cells were measured with a mouse Aβ42 Elisa Kit (27721, IBL, Germany). To measure Aβ42 in brain tissue, 0.05 g of mouse brain tissues were weighed and homogenized using 2ml PBS with a protease inhibitor (cocktail, IKM1020, Solarbio). After centrifugalization at 4 □ for 30 min at 12000 g, the extracts’ supernatants were analyzed using the ELISA method after total protein quantification. To quantify levels of Aβ42 produced from primary neuron cells, the cell supernatants were ultrafiltrated with an Ultrafiltration tube (4-kD Millipore), centrifugalization, and testing. Cell homogenate was prepared in 1ml PBS with cocktail and quantified using the BCA method before being measured by ELISA.

### Immunohistochemistry

Tissue slices were prepared as described before ^42^. The sections were incubated with Aβ (ab201060, Abcam, Cambridge, UK), BACE1 (#5606S, Cell Signaling Technology, Boston, USA), PPARγ (#2435, Cell Signaling Technology, Boston, USA), Notch1 (#3608, Cell Signaling Technology, Boston, USA), p-tau S202 (ab108387, Abcam, Cambridge, UK), p-tau T231(ab151559, Abcam, Cambridge, UK), p-tau S396 (ab109390, Abcam, Cambridge, UK), tau (ab75714, Abcam, Cambridge, UK) antibodies overnight at 4°C and then incubated with Alexa Fluor 488-donkey anti-rabbit IgG (H□+□L) (1:200, Life Technologies, Gaithersburg, MD, USA, #A21208) for 1h, then incubated with a biotin-conjugated secondary antibody for 30□min, followed by incubation with DAB for 10□s and hematoxylin staining for 30□s. The IHC signals were analyzed using ImageJ.

### Cell culture experiments

HT22 cells were purchased from the Cell Bank of the Chinese Academy of Sciences (Shanghai, China). HT22 cells were cultured and grown to confluence in DMEM supplemented with 10% FBS (Gibco BRL), 100 U/mL penicillin, and 100 U/mL streptomycins (Gibco BRL).

### Primary culture of hippocampal neurons

Primary neurons were obtained from the hippocampus of C57/BL6J mice at age 1-3 days. Before culturing, the newborn pup was euthanized and dipped into 70% ethanol for 3 min. First, the infant pup hippocampus was isolated with eye tweezers observed under the stereomicroscope, and excess soft tissue was removed. Second, hippocampal tissue in PBS buffer was cut up with scissors gently and blown with a 1ml pipette until it was not visible. Next, the cell suspension was transferred to a 15ml centrifuge tube and centrifuged at 1000 rpm for 5 min at room temperature. Cell precipitation was suspended and cultured with 2-3mL primary neural stem cell (NSC) suspension (Thermo Fisher Scientific, 21103049) in a 37 °C, 5% CO_2_ cell incubator for 3 days, changing half medium every 2 days. After 7 days, the cell suspension was transferred to a 15ml centrifuge tube, centrifuged, and recultured with neurobasal, 10%FBS, 1:50 B27(Thermo Fisher Scientific, A3582801), and 1:100 bFGF (Thermo Fisher Scientific, #RP-8626). One day later, the medium was changed to neurobasal (2%FBS, 1:50 B27, and 1:100 bFGF), culturing for 21 more days. The immunofluorescence technique was used with the neuron-specific marker (MAP2, #4542, Cell Signaling Technology, Boston, USA) to determine the purity of neurons.

### siRNA, shRNA, and adenovirus transfection

Notch1 siRNA and control siRNA were purchased from RiboBio (Guangzhou, China). HES1 shRNA and control shRNA were purchased from Qingke (Guangzhou, China). Green fluorescent protein-adenovirus (Ad-GFP) and Kallistatin-adenovirus (Ad-KAL) were provided by Dr. Jianxing Ma (University of Oklahoma Health Sciences Center). According to the manufacturer’s instructions, the transfections were performed at approximately 60% confluency using Lipofectamine®3000 transfection reagent (Invitrogen) or RNAiMAX. After 24□h, interference confirmation was conducted using real-time quantitative PCR (RT-qPCR) and Western blot.

### RNA isolation and quantitative RT-PCR

Total RNA extraction, reverse transcription of cDNA, and real-time quantitative PCR were performed as described previously ^43^. BACE1 forward: GGAGCCCTTCTTTGACTCCC; BACE1 reverse: CAATGATCATGCTCCCTCCCA; ADAM9 forward: GGAAGGCTCCCTACTCTCTGA; ADAM9 reverse: CAATTCC-AAAACTGGCATTCTCC; ADAM10 forward: ATGGTGTTGCCGACAGTGTTA; ADAM10 reverse: GTTTGGCACGCTGGTGTTTTT; ADAM17 forward: GGAT-CTACAGTCTGCGACACA; ADAM17 reverse: TGAAAAGCGTTCGGTACTTGAT; β-actin forward: GCACTCTTCCAGCTTCCTT; β-actin reverse: GTTGGCGTACAG-GTCTTTGC.

### Western blot

Western blot was performed as described previously ^40, 43^. Equal amounts of protein were subjected to western blot analysis. Blots were probed with antibodies against Kallistatin (ab187656, Abcam, Cambridge, UK), Aβ (ab201060, Abcam, Cambridge, UK), Presenilin-1 (ab76083, Abcam, Cambridge, UK), BACE1 (#5606S, Cell Signaling Technology, Boston, USA), APP (#2452S, Cell Signaling Technology, Boston, USA), MAP2 (#4542, Cell Signaling Technology, Boston, USA), PPARγ (#2435, Cell Signaling Technology, Boston, USA), SP1 (#9389, Cell Signaling Technology, Boston, USA), YY1 (#46395, Cell Signaling Technology, Boston, USA), Notch1 (#3608, Cell Signaling Technology, Boston, USA), Hes1 (#11988, Cell Signaling Technology, Boston, USA), p-tau S202 (ab108387, Abcam, Cambridge, UK), p-tau T231(ab151559, Abcam, Cambridge, UK), p-tau S396 (ab109390, Abcam, Cambridge, UK), tau (ab75714, Abcam, Cambridge, UK), GSK3β(#70109S, Cell Signaling Technology, Boston, USA), p-GSK3β Ser9 (#9323, Cell Signaling Technology, Boston, USA), β-actin (A5441-2ml, Sigma, CA, USA), Caveolin-1(SZ02-01, Huabio, China), GAPDH (200306-7E4, Zen-bio, China), anti-Mouse (#PI200, Vector Laboratories, Burlingame, CA, USA), anti-Rabbit (#PI1000, Vector Laboratories, Burlingame, CA, USA). The signal intensity was quantified using ImageJ (NIH).

### Statistical Analysis

The results are expressed as mean ± SD. Student’s *t*-test was applied for comparisons of parametric data between two groups, and one-way ANOVA followed by LSD *t*-test was used to compare differences between more than two different groups (GraphPad Prism software). A *P* value less than 0.05 was considered statistical significance.

## List of abbreviations

Aβ: amyloid β
p-tau: hyperphosphorylated tau
AD: Alzheimer’s disease
T2DM: type 2 diabetes mellitus
FBG: fasting blood glucose
TG: triglyceride
KAL-TG: Kallistatin-transgenic
WT: wild type mice
APP: amyloid precursor protein
BACE1: β-site APP cleaving enzyme 1
BMI: body mass index
ICD-10: The International Statistical Classification of Diseases and Related Health Problems 10th Revision
NINCDS-ADRDA: the National Institute of Neurological and Communicative Disorders and Stroke and the Alzheimer’s Disease and Related Disorders Association
MMSE: mini-Mental State Examination
KAL-TG-RSG: rosiglitazone-treated group
KAL-TG-Feno: fenofibrate-treated group
MWM: morris water maze
RT-qPCR: real-time quantitative PCR

## Declarations

## Consent for publication

Not applicable.

## Availability of data and materials

All the data supporting the conclusions of the current study are presented in the figures and they are available from the corresponding authors upon reasonable request. There are no restrictions on data availability. Source data are provided with this paper.

## Acknowledgements

We are thankful to Zhongshan City People’s Hospital, Zhongshan Third People’s Hospital, Guangdong Provincial People’s Hospital, and Sun Yat-sen Memorial Hospital for kindly providing serum samples for the analyses of this manuscript. We thank Professor Boxing Li for his valuable advice on our study.

## Authors’ contributions

G. Gao, X Yang, B Jiang, and P Jiang were involved in the concept and design of the study. W. Qi, Y. Long, Z. Li, Z. Zhao J. Shi, D Zhu, Z Zhao, W. Xie, L. Wang, T Zhou, Mingting Liang were responsible for conducting the experiments. W. Qi, Y. Long and T Zhou drafted the manuscript and G. Gao revised the manuscript. P Jiang, Y. Long and Z. Li were responsible for data analysis. All authors contributed to the interpretation of data and provided revisions to the manuscript. G. Gao will act as guarantor for the study. All authors read and approved the final manuscript. W. Qi, Y. Long and Z. Li contributed equally to this study.

---

**Table. S1.**
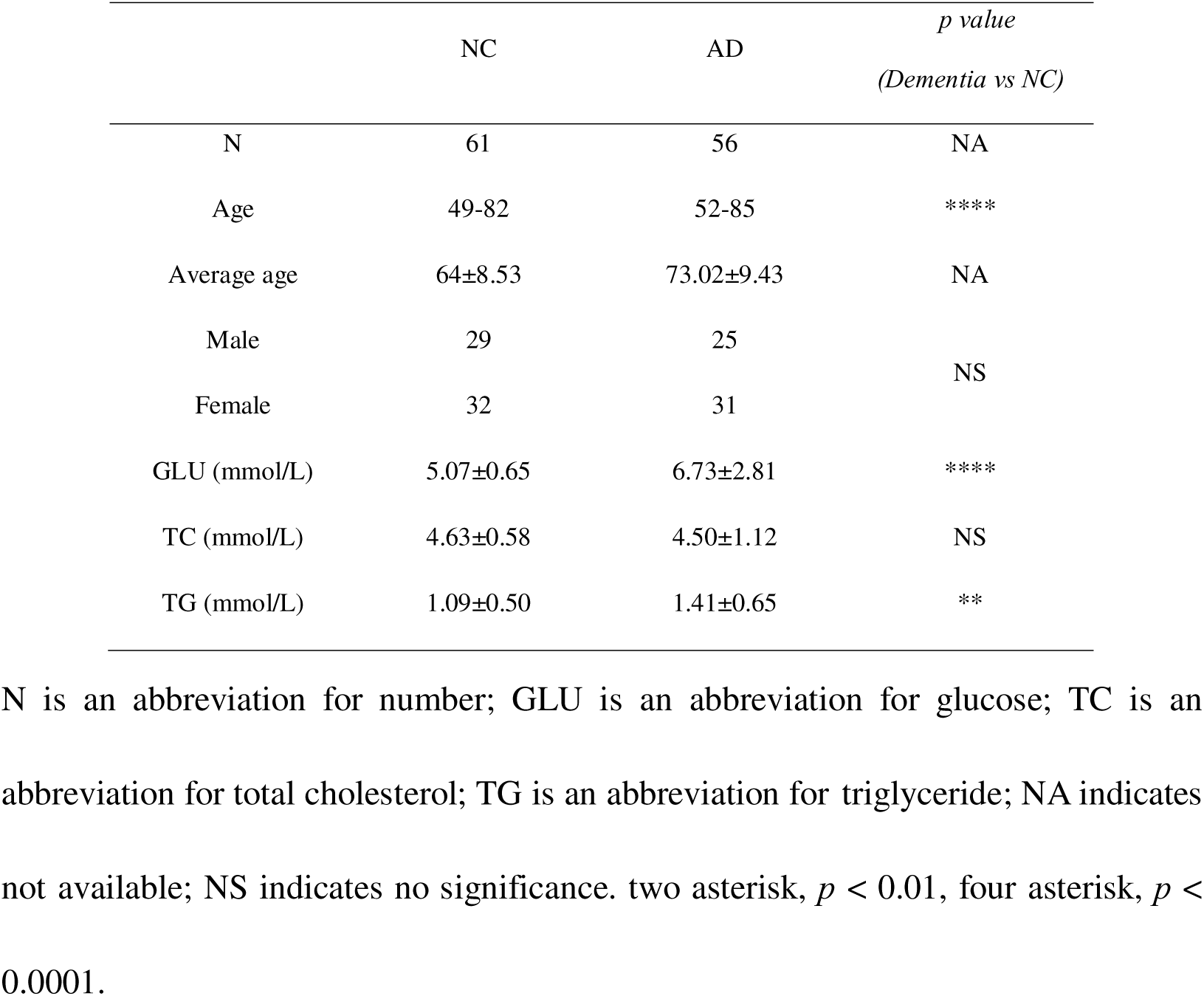
Clinical characteristic of AD patients.

|  | NC | AD | *p value*<br>(*Dementia vs NC*) |
| --- | --- | --- | --- |
| N | 61 | 56 | NA |
| Age | 49-82 | 52-85 | **** |
| Average age | 64±8.53 | 73.02±9.43 | NA |
| Male | 29 | 25 | NS |
| Female | 32 | 31 |  |
| GLU (mmol/L) | 5.07±0.65 | 6.73±2.81 | **** |
| TC (mmol/L) | 4.63±0.58 | 4.50±1.12 | NS |
| TG (mmol/L) | 1.09±0.50 | 1.41±0.65 | ** |

N is an abbreviation for number; GLU is an abbreviation for glucose; TC is an abbreviation for total cholesterol; TG is an abbreviation for triglyceride; NA indicates not available; NS indicates no significance. two asterisk, $p < 0.01$ , four asterisk, $p <$ 0.0001.

**Table. S2.**
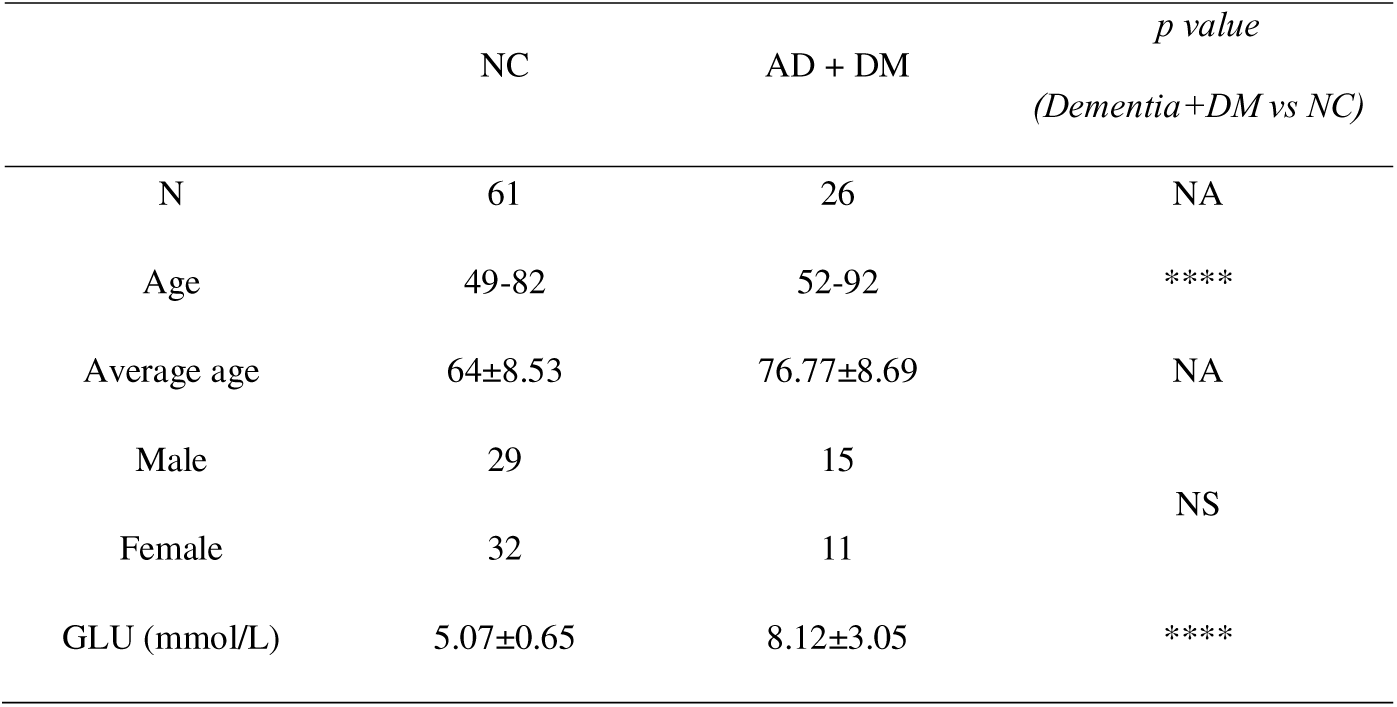
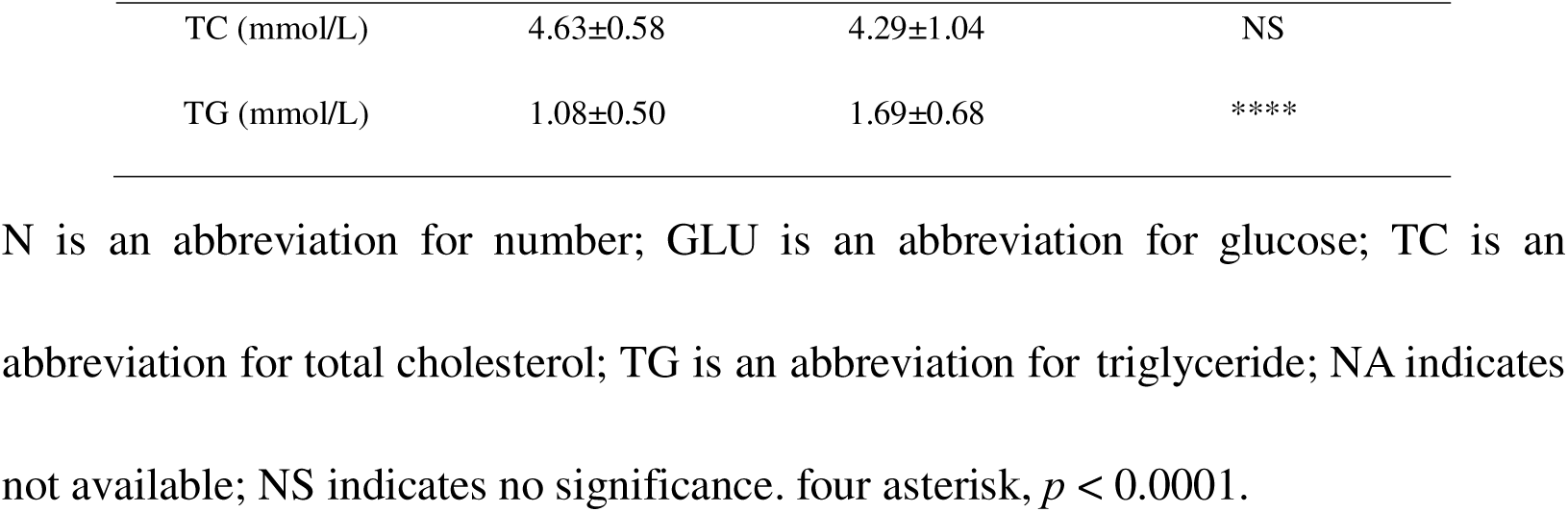
Clinical characteristic of AD patients with DM.

**Fig. S1.**
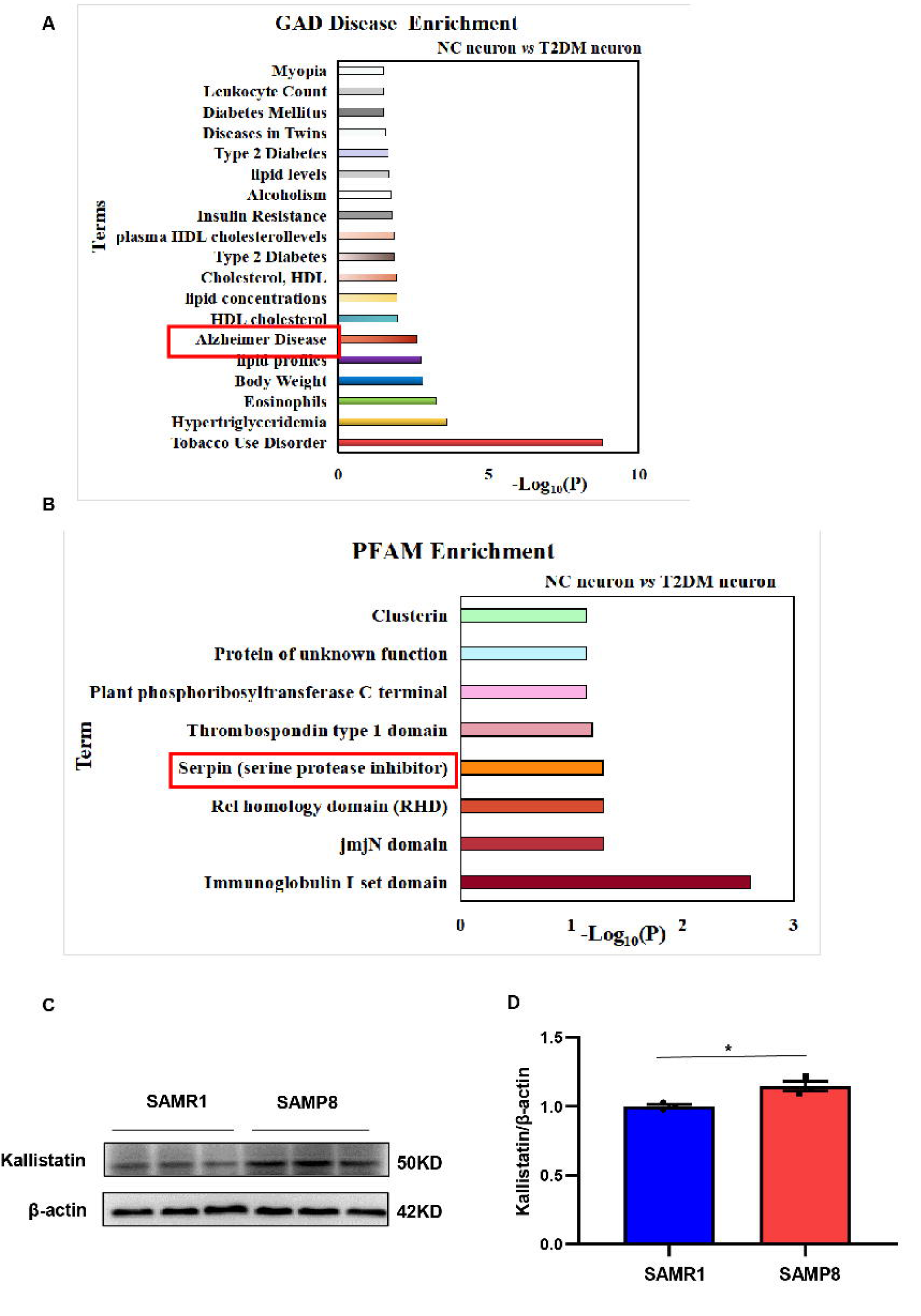
(A-B) GAD disease enrichment analysis (A) and PFAM analysis (B) result. Differentially expressed genes in neurons were obtained from GSE161355, and GAD disease enrichment was analyzed on David database. (C-D) Western blot analysis of Kallistatin expression in aging model SAMP8 and corresponding control SAMR1 mice hippocampal tissue samples, then statistically analyzing the above results. β-Actin served as a loading control. Error bars represent the standard deviation (SD); one asterisks, *p* < 0.05.

**Fig. S2.**
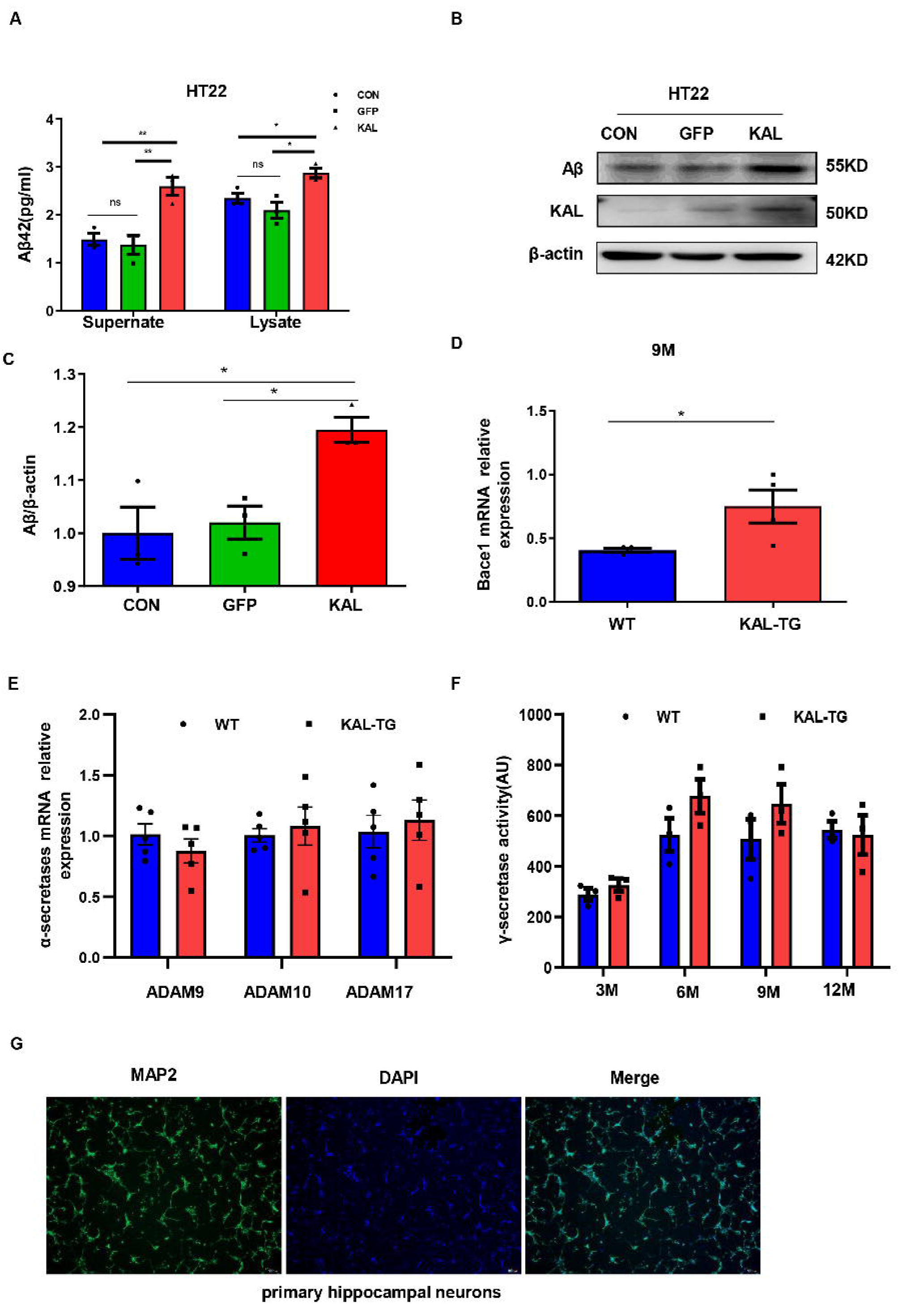
(A) HT22 cells were infected with adenovirus to overexpress Kallistatin for 48h. Aβ42 concentration of supernate and cell lysate was quantified by ELISA. (B-C) Western blot analysis of Aβ protein level in HT22 cells infected with overexpressing Kallistatin adenovirus and control groups for 48h, then statistical analysis of Kallistatin protein levels. (D) BACE1 mRNA expression in hippocampal tissue. (E) α-secretases (ADAM9, ADAM10, ADAM17) mRNA expression in hippocampal tissue. (F) γ-secretase activity of each group’s hippocampal tissue was measured by ELISA. (G) Primary hippocampal neurons were identified with MAP2. Error bars represent the standard deviation (SD), one asterisk, *p* < 0.05, two asterisks, *p* < 0.01.

**Fig. S3.**
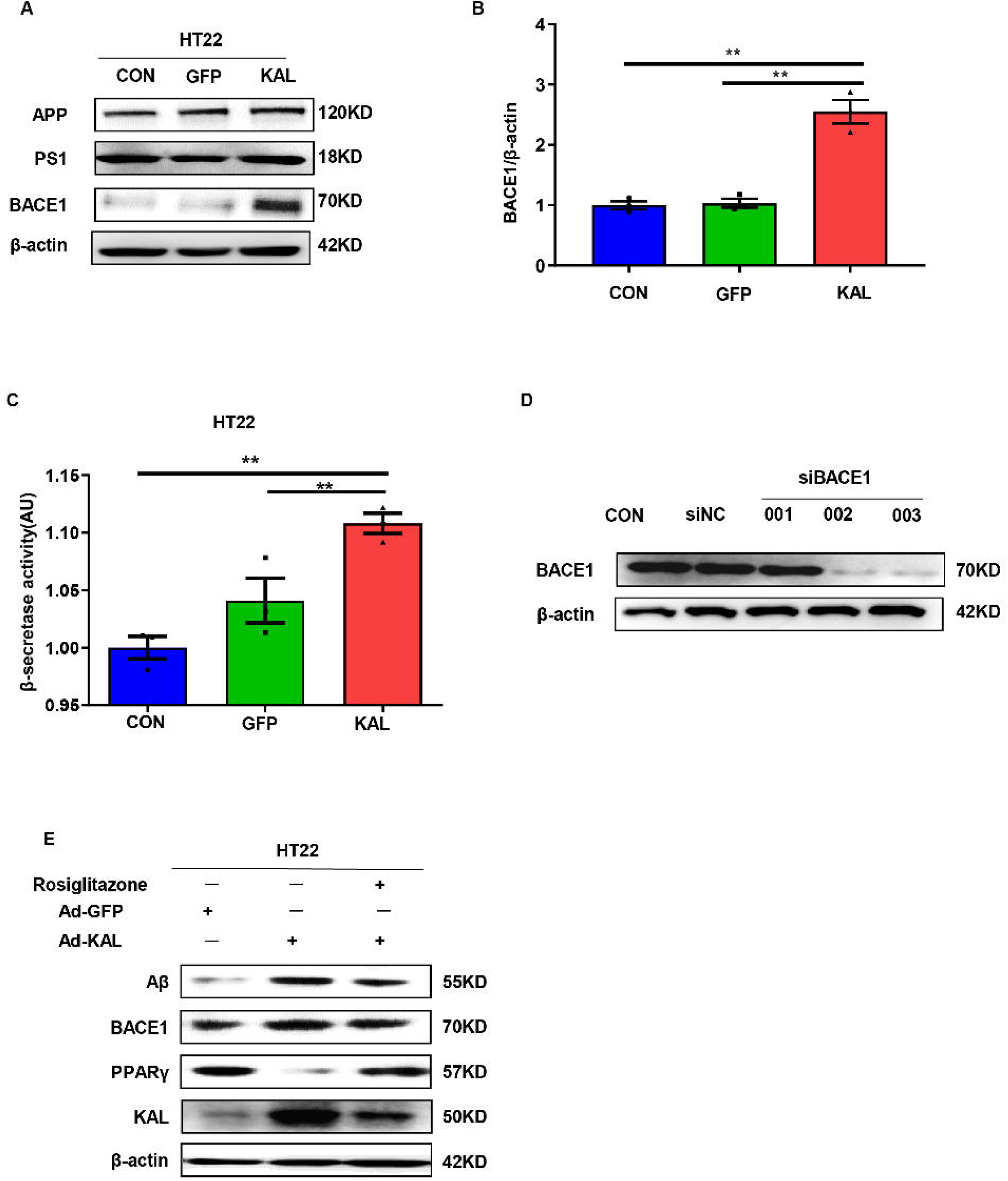
(A) The relevant protein levels in HT22 cells infected with overexpressing Kallistatin adenovirus during Aβ generation were determined by western blot analysis. (B) Statistical analysis of BACE1 expression in HT22 cells. (C) β-secretase activity of HT22 cells infected with overexpressing Kallistatin adenovirus and control adenovirus was measured by ELISA. (D) Primary hippocampal neurons were infected with BACE1 siRNA for 72h. Western blot analysis of BACE1 protein levels. (E) HT22 cells were treated with PPARγ agonist rosiglitazone (10nM) for 12h, then infected with adenovirus to overexpress Kallistatin for 48h. Western blot analysis of Aβ and BACE1 protein levels. β-actin served as a loading control. Error bars represent the standard deviation (SD), two asterisks, *p* < 0.01.

**Fig. S4.**
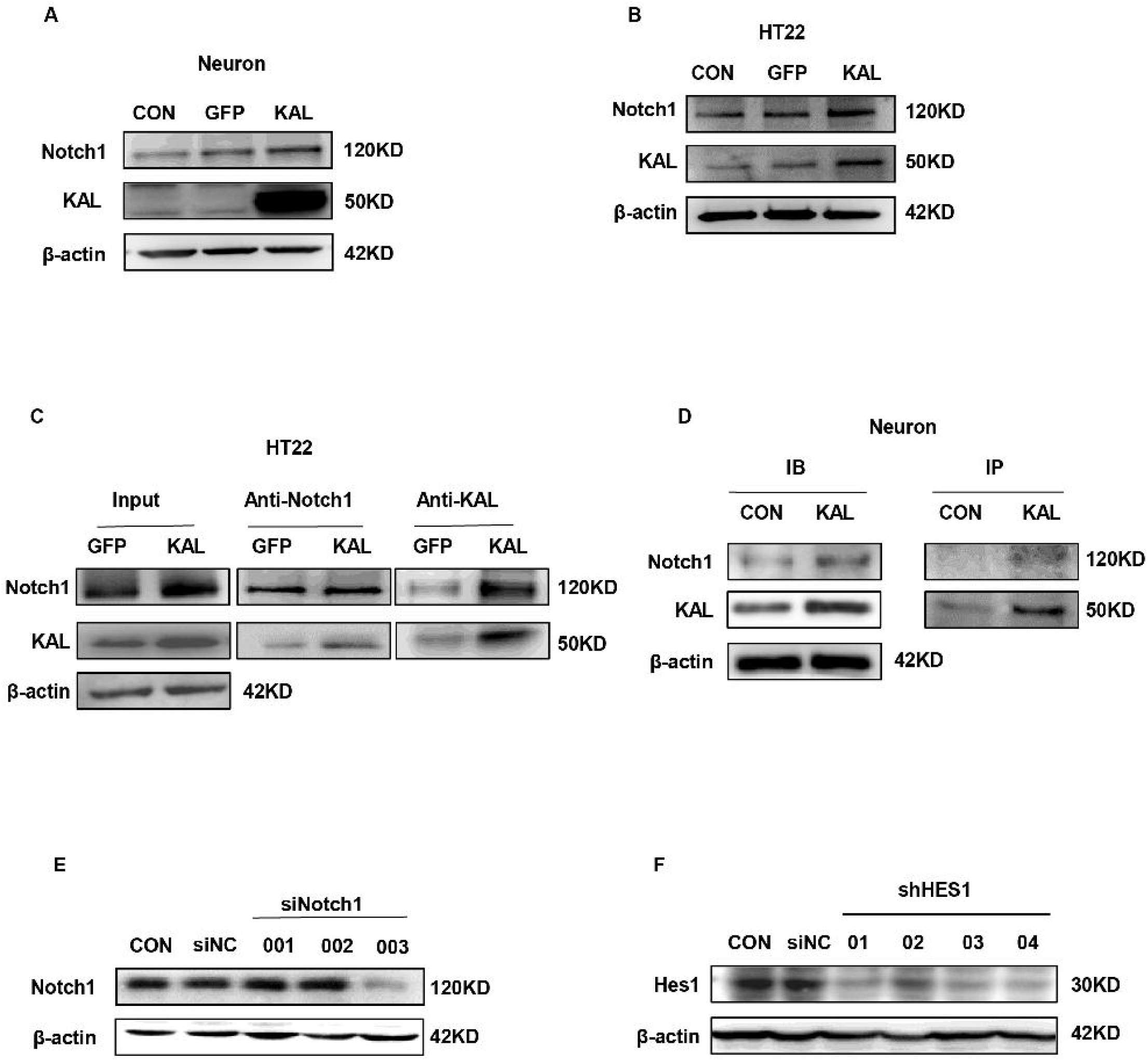
(A-B) Western blot analysis of Notch1 protein level in primary hippocampal neurons and HT22 cells infected with overexpressing Kallistatin adenovirus and control groups. (C) HT22 cells were infected with overexpressing Kallistatin adenovirus for 48h, and Co-IP analysis was conducted to verify whether Kallistatin can bind to the Notch1 receptor. β-actin served as a loading control. (D) Primary hippocampal neurons were treated with Kallistatin protein for 72h, then IP analysis. (E) Primary hippocampal neurons were infected with Notch1 siRNA for 72h. Western blot analysis of BACE1 protein levels. (F) HT22 cells were infected with HES1 shRNA for 48h. Western blot analysis of HES1 protein levels.

**Fig. S5.**
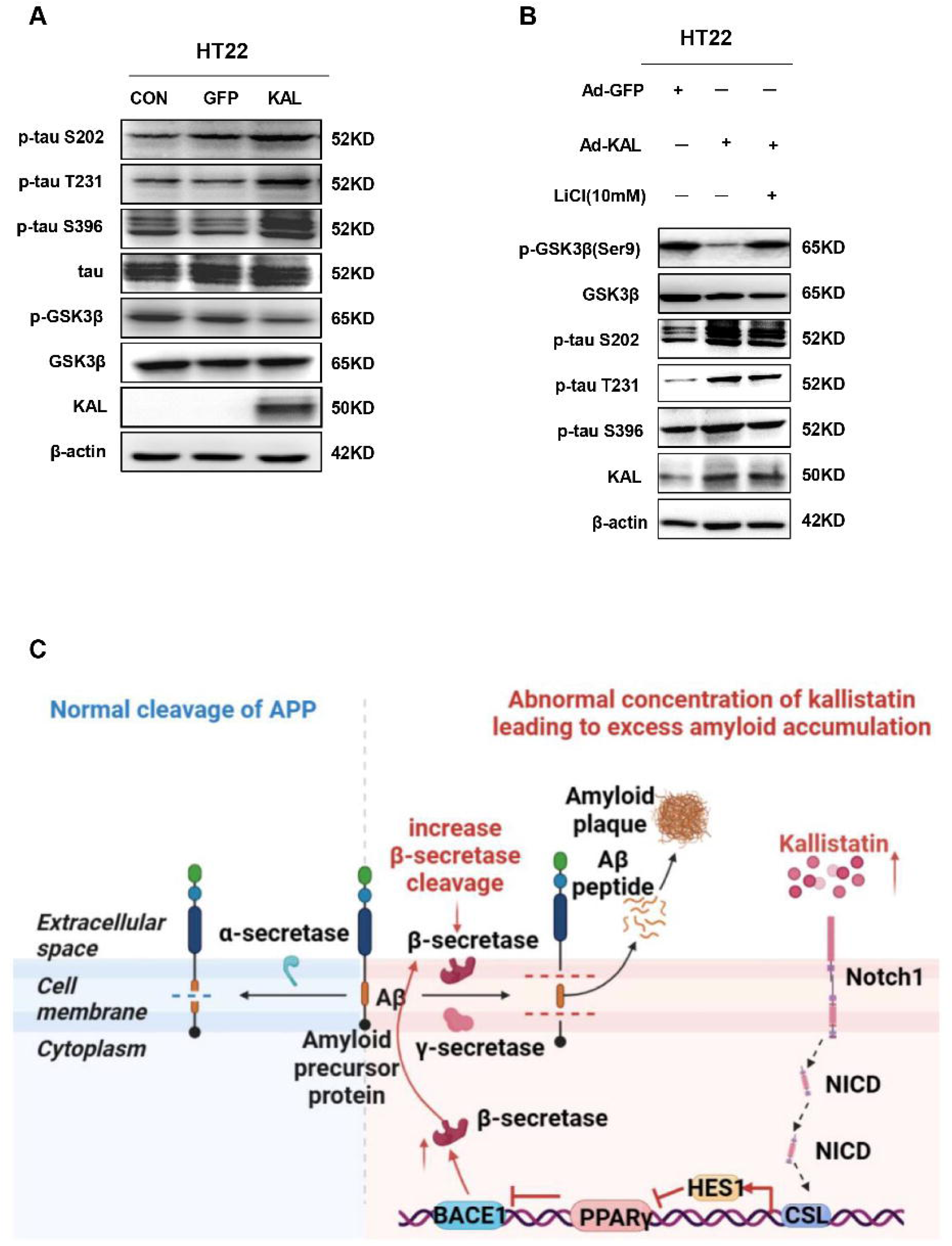
(A) Western blot analysis of GSK-3β, p-GSK-3β, tau, p-tau(Ser9, T231, S396) in HT22 cells infected with overexpressing Kallistatin adenovirus and control groups. (B) Western blot analysis of GSK-3β, p-GSK-3β, p-tau(Ser9, T231, S396) in HT22 cells infected with overexpressing Kallistatin adenovirus and control groups for 24h, then treated with LiCl(10 mM) for 24h. (C) Simplified model depicting the pathway of Aβ regulated by Kallistatin.

